# Intracranial recordings from human auditory cortex reveal a neural population selective for song

**DOI:** 10.1101/696161

**Authors:** Sam V Norman-Haignere, Jenelle Feather, Dana Boebinger, Peter Brunner, Anthony Ritaccio, Josh H McDermott, Gerwin Schalk, Nancy Kanwisher

## Abstract

How are neural representations of music organized in the human brain? While neuroimaging has suggested some segregation between responses to music and other sounds, it remains unclear whether finer-grained organization exists within the domain of music. To address this question, we measured cortical responses to natural sounds using intracranial recordings from human patients and inferred canonical response components using a data-driven decomposition algorithm. The inferred components replicated many prior findings including distinct neural selectivity for speech and music. Our key novel finding is that one component responded nearly exclusively to music with singing. Song selectivity was not explainable by standard acoustic features and was co-located with speech- and music-selective responses in the middle and anterior superior temporal gyrus. These results suggest that neural representations of music are fractionated into subpopulations selective for different types of music, at least one of which is specialized for the analysis of song.

---

Music is a quintessentially human capacity that is present in some form in nearly every society (Savage et al., 2015; Lomax, 2017; Mehr et al., 2018), and that differs substantially from its closest analogues in non-human animals (Patel, 2019). Researchers have long debated whether the human brain has neural mechanisms dedicated to music, and if so, what computations those mechanisms perform (Patel, 2012; Peretz et al., 2015). These questions have important implications for understanding the organization of auditory cortex (Leaver and Rauschecker, 2010; Norman-Haignere et al., 2015), the neural basis of sensory deficits such as amusia (Peterson and Pennington, 2015; Norman-Haignere et al., 2016; Peretz, 2016), the consequences of auditory expertise (Herholz and Zatorre, 2012), and the computational underpinnings of auditory behavior (Casey, 2017; Kell et al., 2018).

Neuroimaging studies have suggested that representations of music diverge from those of other sound categories in non-primary human auditory cortex: some non-primary voxels show partial selectivity for music compared with other categories (Leaver and Rauschecker, 2010; Fedorenko et al., 2012; Angulo-Perkins et al., 2014), and recent studies from our lab, which modeled voxels as weighted sums of multiple response profiles, inferred a component of the fMRI response with clear selectivity for music (Norman-Haignere et al., 2015; Boebinger et al., 2020) that was distinct from nearby speech-selective responses. However, little is known about how neural responses with the domain of music are organized, such as whether distinct subpopulations exist that are selective for particular types or features of music (Casey, 2017). As a consequence, we know little about the neural code for music.

Here, we tested for finer-grained selectivity within the domain of music by using intracranial recordings from the human brain (electrocorticography or ECoG), which have better spatiotemporal resolution than any other human neuroscience method. We measured ECoG responses to a diverse set of 165 natural sounds (**Fig 1A**), and submitted these responses to a novel decomposition method that is well-suited to the statistical structure of ECoG to reveal dominant response components of auditory cortex. The components revealed by this analysis replicated many prior findings including tonotopic frequency selectivity (Humphries et al., 2010; Costa et al., 2011; Moerel et al., 2012; Baumann et al., 2013), spectrotemporal modulation tuning (Schönwiesner and Zatorre, 2009; Barton et al., 2012; Santoro et al., 2014), spatially organized onset responses (Hamilton et al., 2018), as well as selectivity for speech, music, and vocalizations (Belin et al., 2000; Leaver and Rauschecker, 2010; Angulo-Perkins et al., 2014; Overath et al., 2015). Our key novel finding is that one of these components responded nearly exclusively to music with singing. This finding provides clear evidence that the human brain contains neural populations specific to the analysis of song.

**Figure 1.**
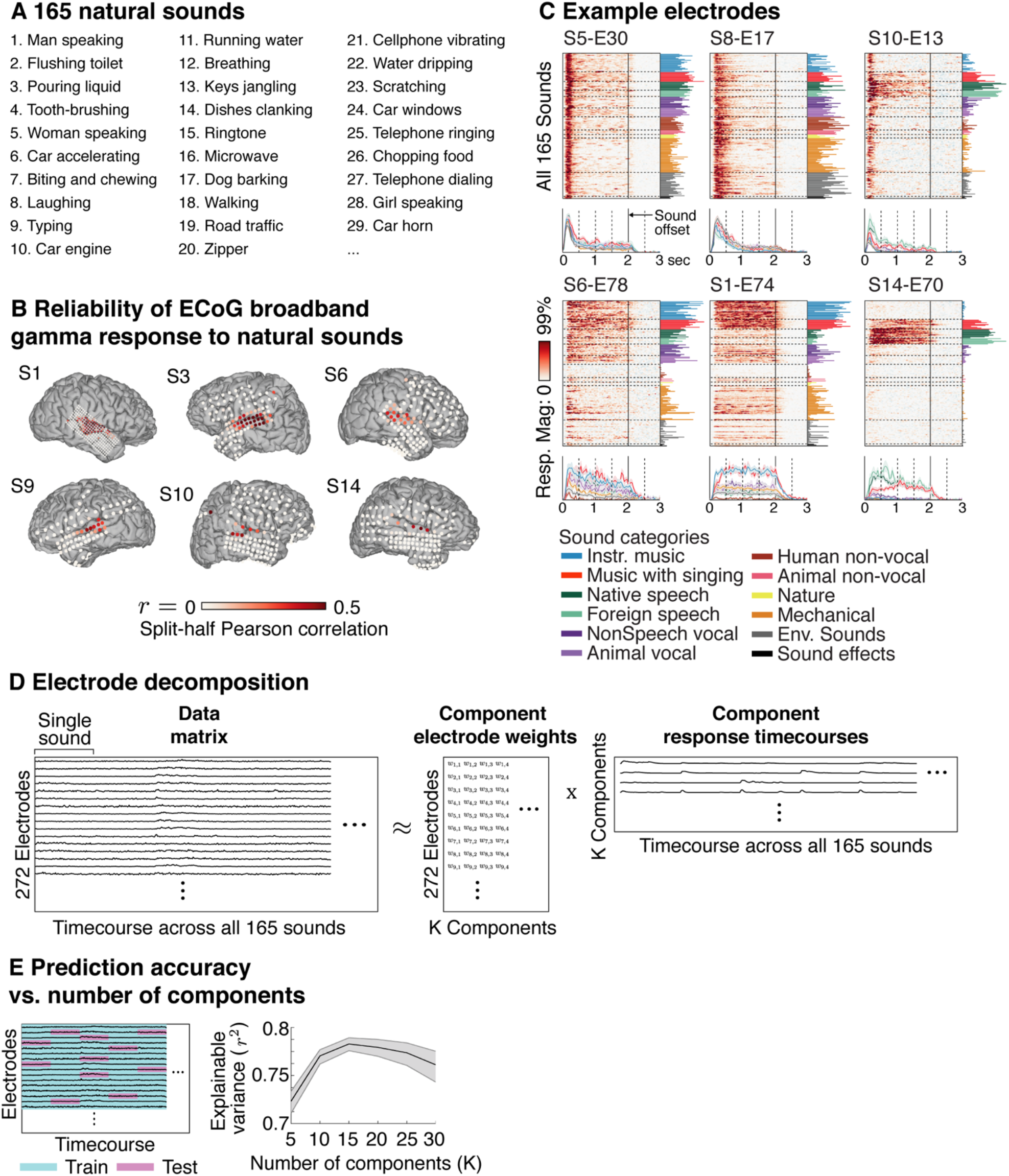
Overview of experiment and decomposition method. **A**, The sound set was composed of 165 commonly heard sounds, each 2-seconds in duration (Norman-Haignere et al., 2015). **B**, Electrodes were selected based on the split-half-reliability of their broadband gamma response (70-140 Hz) to natural sounds. This panel plots maps of the split-half correlation for six example subjects (ordered based on the total number of reliable electrodes). **C**, The broadband gamma response timecourse of several example electrodes to all 165 sounds, plotted as a raster. The time-averaged response to each sound is plotted to the right of the raster. The sounds have been grouped and colored based on membership in one of 12 sound categories (determined primarily based on subject ratings; see *Sound Category Assignments* in Methods). Below each raster, we plot the average response to each category with greater than 5 exemplars. Error bars plot the median and central 68% of the sampling distribution, computed via bootstrapping across sounds. We plot the most reliable electrode for several example subjects to illustrate the diversity of responses. **D**, The data were represented as a matrix, where each row contains the full response timecourse of each electrode (from 0 to 3 seconds post-stimulus onset), concatenated across all 165 sounds tested. The data matrix was approximated as the product of a response timecourse matrix, which contains a small number of canonical response timecourses that are shared across all electrodes, with an electrode weight matrix that expresses the contribution of each component timecourse to each electrode. **E**, Cross-validation was used to compare models (**Fig S2C**) and determine the number of components. The data matrix was divided into cells, with one cell containing the response timecourse of a single electrode to a single sound. The model was trained on a randomly chosen subset of 80% of cells, and was then asked to predict responses for the remaining 20% of cells. This plot shows the squared test correlation between the measured and predicted response (averaged across all electrodes) as a function of the number of components. The correlation has been noise-corrected using the test-retest reliability of the electrode responses so that it provides a measure of explainable variance. Error bars plot the median and central 68% of the sampling distribution (equivalent to 1 standard error for a Gaussian), computed via bootstrapping across subjects.

## Results

### Intracranial recordings

We identified a set of 272 electrodes across 15 patients with reliable broadband gamma (70-140 Hz) power responses to the sound set (split-half correlation > 0.2; **Fig 1B** plots the split-half correlation for all electrodes). We focused on broadband gamma, because it is thought to reflect aggregate spiking in a local region (Steinschneider et al., 2008; Whittingstall and Logothetis, 2009; Ray and Maunsell, 2011). Sound-responsive electrodes were nearly always located on or near the superior temporal gyrus (STG), as expected. The number of reliable electrodes varied substantially across subjects due to the sparse, clinically driven coverage of ECoG grids (**Fig 1B**).

To illustrate the format of the data and the heterogeneity of the electrode responses, we plot the most reliable electrode from several example subjects in **Figure 1C**. We plot the response timecourse of each electrode to all 165 sounds as a stack of raster plots, along with the time-averaged response to each sound (to the right of the raster) and the average response timecourse to several different sound categories (below the raster) (see *Sound Category Assignments* in Methods). Electrodes exhibited a variety of responses including strong responses at sound onset (e.g. S5-E30) and selective responses to speech (e.g. S14-E70).

### Electrode decomposition

Rather than analyze individual electrodes, we attempted to identify a small number of canonical response timecourses across the sound set (components) that could collectively explain most of the response variance across all 272 electrodes when weighted and summed. Each component timecourse could potentially reflect a different neuronal subpopulation in auditory cortex, with the weights providing an estimate for the contribution of each subpopulation to each electrode.

To identify components, we represented the electrode responses as a matrix, in which each row contained the concatenated response timecourses of a single electrode to all 165 sounds (**Fig 1D**). We approximated this matrix as the product of a component response timecourse matrix and a component electrode weight matrix. In general, the problem of matrix factorization – finding a set of response components whose weighted sum best explains the data – is ill-posed and needs to be constrained by additional statistical criteria. We identified three statistical properties of auditory broadband gamma activity that are relevant to component modeling (**Fig S1**): (1) relative to silence, sound-driven responses are nearly always excitatory (suppressive responses accounted for <1% of the response power); (2) responses are sparse across both time/stimuli and electrodes; (3) responses are temporally smooth, but the extent of this smoothness varies across electrodes. We designed a model that captured all of these statistical properties by convolving a set of sparse/non-negative components with a learned smoothing kernel (**Fig S2**; see Methods for details). We focus on the results of this model because it yielded better cross-validated prediction accuracy than competing models (**Fig S2C**). But we note that our key results were evident using a simpler model that only imposed non-negativity on the responses and weights (**Fig S3**).

Using a standard cross-validation procedure, in which we trained and tested on non-overlapping pairs of sounds/electrodes, we found that we could estimate ~15 components before overfitting (**Fig 1E**). We focus on a subset of 10 reliable components that were also present in a simpler NMF model (**Fig S3**), were stable across the number of model components (**Fig S4**), and explained responses across multiple subjects (**Fig S5**; **Fig S6** plots the 5 less reliable components from a 15-component model).

### Speech and music-selective components

We first describe three components that responded selectively to speech or music (**Fig 2**), even though categories played no role in the decomposition algorithm. For each component, we plot its response (**Fig 2A**) as well as an anatomical map of its electrode weights (**Fig 2B**) (note that anatomy also played no role in the decomposition).

**Figure 2.**
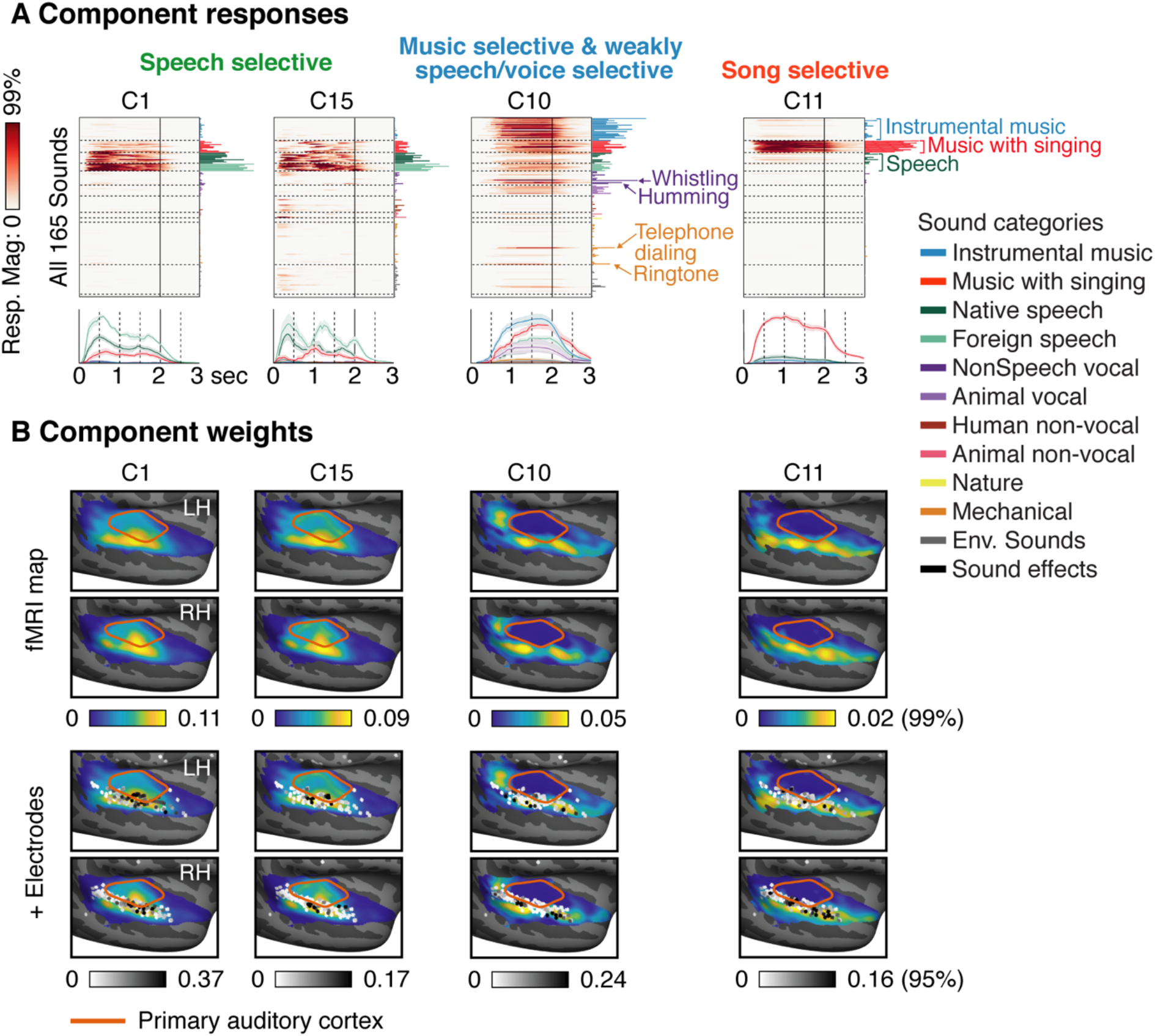
Category-selective components. The response and anatomy of four components that responded nearly exclusively to either speech or music. **A**, The response timecourse of each component. Same format as the example electrodes shown in Figure 1C. **B**, Anatomical distribution of each component, measured using fMRI (top panel) and ECoG (bottom panel; darker dots indicate greater electrode weights). fMRI maps are shown in both panels to facilitate comparison. See text for details of how fMRI maps were computed.

Because ECoG coverage is highly restricted, we complemented the electrode weight map with a second anatomical map, computed using a large dataset of fMRI responses to the same sound set across 30 subjects from two studies (~176 hours of scanning) (Norman-Haignere et al., 2015; Boebinger et al., 2020). This map was computed by projecting the response of each fMRI voxel onto the time-averaged component responses inferred using ECoG, and then averaging across all 30 subjects. This method enables us to leverage the dense and comprehensive coverage of fMRI data to provide an estimate of the full weight map for each ECoG-derived component. As expected, the split-half correlation of the fMRI weight maps across subjects was much higher than the split-half correlation of the ECoG maps due to superior coverage and more subjects (**Fig S7**). Moreover, the correlation between fMRI and ECoG maps for corresponding components was slightly higher than the split-half correlation of the ECoG maps themselves, and much higher than that for mismatching components (p < 0.001 via bootstrapping across the fMRI subjects) (**Fig S7**). These findings suggest a close correspondence between the fMRI and ECoG maps that is primarily limited by the sparse coverage of ECoG recordings, and thus demonstrates the utility of combining the precision of ECoG recordings with the spatial coverage of fMRI. We used the fMRI weight maps to test for effects of laterality, since we had dense, bilateral coverage from a large number of subjects (**Fig S8** shows the average difference in weights between the left and right hemisphere).

Two components (C1, C15) responded nearly exclusively to speech, producing little to no response for all other sounds including non-speech vocalizations. The response in these components was similar to native and foreign speech sounds, consistent with prior work showing that speech selectivity in STG is not driven by linguistic meaning (Norman-Haignere et al., 2015; Overath et al., 2015). C1 and C15 responded at different moments within each speech utterance, plausibly driven by previously reported selectivity for speech features such as phoneme classes (e.g. Mesgarani et al., 2014), but the acoustic features which drove this variation were difficult to characterize because of the relatively small amount of speech in our corpus. The weights for these speech-selective components were primarily clustered in middle STG, with no significant difference between the two hemispheres (p > 0.73 uncorrected for the number of components via bootstrapping across subjects), again consistent with prior work (Leaver and Rauschecker, 2010; Mesgarani et al., 2014; Overath et al., 2015; Boebinger et al., 2020). We note that the time-averaged response of C1 and C15 was very similar, which limited our ability to spatially distinguish these two components with fMRI.

One component (C10) responded strongly to both instrumental music and music with singing (average[instrumental music, sung music] > average[all non-music categories]: p < 0.001 via bootstrapping, Bonferroni-corrected for the number of components). This component produced an intermediate response to speech, suggesting that music and speech were not perfectly disentangled by our component analysis, perhaps due to limited coverage of the lateral sulcus where music selectivity is prominent and speech-selectivity is weak (Norman-Haignere et al., 2015; Boebinger et al., 2020). All other non-music and non-speech sounds produced weak responses. C10 also showed the most delayed response of all of the inferred components (708 ms; measured as the time-to-half maximum of the first principal component of the time x sound response matrix for each component), perhaps suggesting a longer integration window (Norman-Haignere et al., 2020). The weights for this component showed three hotspots in posterior, middle and anterior STG. This anatomical profile is similar to the music-selective component we previously inferred from fMRI data alone (Norman-Haignere et al., 2015; Boebinger et al., 2020), but the cluster in middle STG was more prominent, likely due to the weak speech responses in this component.

### Song selectivity

Our key novel finding is that one component (C11) responded nearly exclusively to sung music: every music stimulus with singing produced a high response and all other sounds, including both speech and instrumental music, produced little to no response. As a consequence, the response to sung music was substantially and significantly higher than the sum of the response to speech and instrumental music, indicating a nonlinear response preference for sung music (sung music > max[English speech, foreign speech] + instrumental music: p < 0.001 via bootstrapping, Bonferroni-corrected). Moreover, the response of C11 could not be explained as a linear combination of our previously discovered fMRI components which showed clear selectively for music and speech individually (**Fig S9**). This finding of nonlinear song selectivity is strengthened by the fact that our decomposition method explicitly models each electrode as a weighted sum of multiple components, and thus if song selectivity simply reflected a sum of speech and music selectivity, the model should not have needed a separate component selective for just sung music. C11 had anatomical weights that clustered exclusively in non-primary auditory cortex, overlapping both speech-selective regions in the middle STG and music-selective regions in anterior STG. C9 also showed a relatively delayed response (298 ms) relative to many of the other components, consistent with its anatomical location in non-primary auditory cortex.

A subtle but important fact about our decomposition algorithm is that the inferred components are not guaranteed to be a linear function of the data (i.e. a weighted sum of the electrode responses) even though the data are approximated as a linear function of the components. This fact is notable because super-additive selectivity for singing cannot be explained by a linear function of speech and music selectivity, and thus the existence of a linear component with super-additive song selectivity would provide direct evidence for nonlinear tuning. To test if such a component existed, we attempted to learn a weighted sum of the electrode responses that yielded a binary song-selective response, using cross-validated regression. We found that this analysis was successful, and yielded a component that was nearly identical to the song-selective component inferred by our decomposition algorithm (**Fig S10**), even in subtle characteristics not explicitly constrained by our regression analysis (i.e. the temporal profile of responses to each sung music stimulus). We note that unlike our decomposition algorithm, this analysis does not depend on statistical assumptions like non-negativity or sparsity, and thus provides complementary evidence for nonlinear song selectivity.

### Selectivity for spectrotemporal modulation statistics

Can speech, music and song selectivity be explained by generic acoustic representations, such as spectrotemporal modulations that appear to drive much of the functional organization of human primary auditory cortex (Schönwiesner and Zatorre, 2009; Barton et al., 2012; Santoro et al., 2014)? This question is relevant since speech and music are known to have distinctive modulation statistics (Singh and Theunissen, 2003; Ding et al., 2017; Elhilali, 2019). We recently introduced an algorithm for synthesizing sounds that are matched to natural sounds in their spectrotemporal modulation statistics, despite being acoustically distinct (**Fig 3A**) (Norman-Haignere and McDermott, 2018). We previously found that fMRI voxels in primary auditory cortex respond similarly to natural and modulation-matched synthetics, but that non-primary regions produced weak responses to the synthetic sounds, presumably because they lack higher-order structure necessary to drive neurons in non-primary regions.

**Figure 3.**
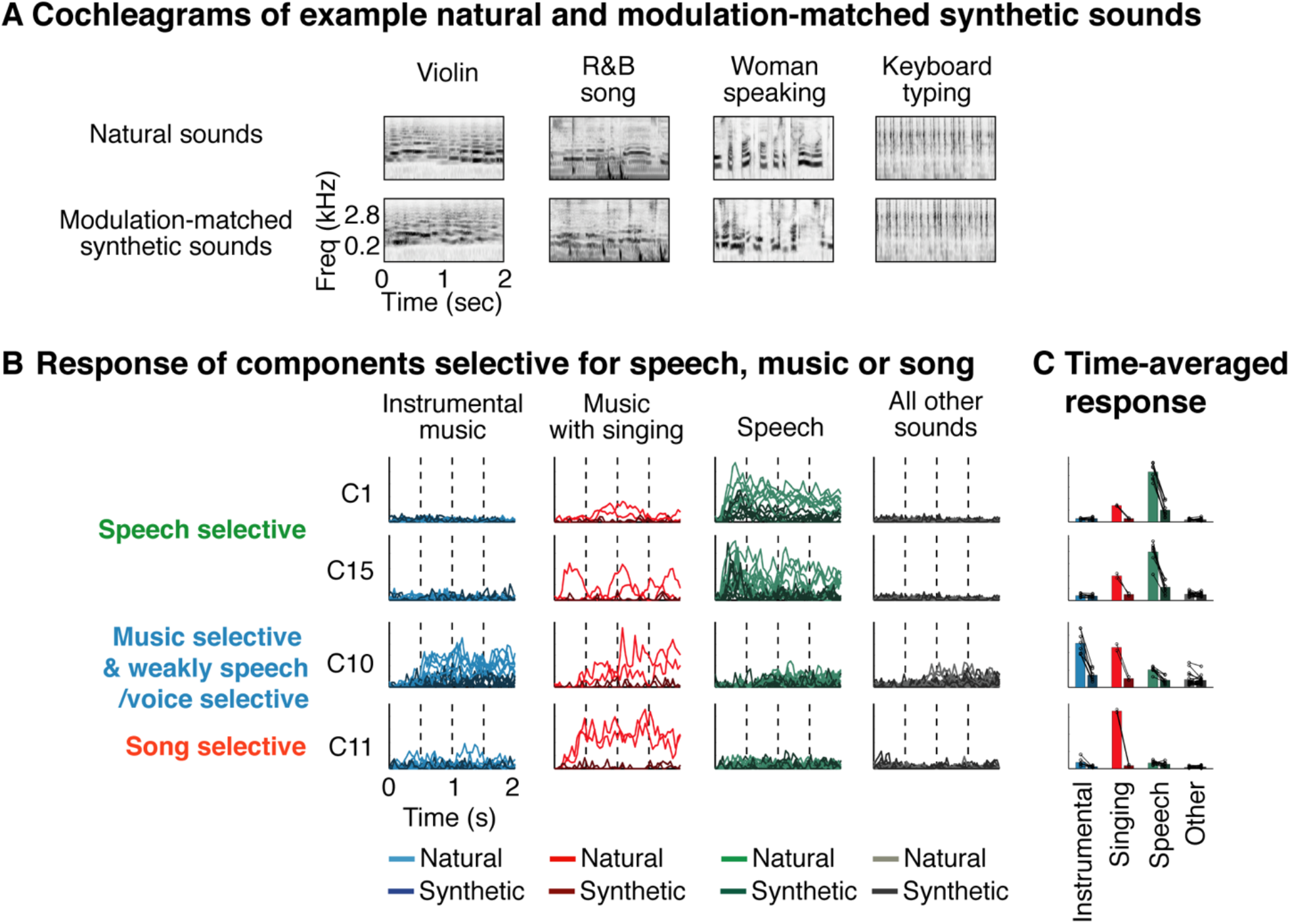
Component responses to natural and modulation-matched synthetic sounds. **A**, Cochleagrams of example natural and corresponding synthetic sounds with matched spectrotemporal modulation statistics (Norman-Haignere and McDermott, 2018). Cochleagrams plot energy as a function of time and frequency, similar to a spectrogram, but measured from filters designed to mimic cochlear frequency tuning. Stimuli lasted 4 seconds, but just the first 2 seconds of the cochleagram is plotted. **B**, The response of the speech, music, and song-selective components, identified in the 165-natural sound experiment, to the natural and modulation-matched sounds of the control experiment. We plot the response timecourse (first 2-seconds) of each component to each natural (lighter colors) and modulation-matched synthetic sound (darker colors). The sounds have been grouped into four categories: instrumental music (blue), music with singing (red), speech (green, both English and foreign), and all other sounds (black/gray). **C**, The time-averaged component response to each pair of natural and modulation-matched sounds (connected circles indicate pairs), along with the mean component response across the natural (lighter bars) and modulation-matched (darker bars) sounds from each category.

We measured responses in a subset of 10 patients to a new set of 36 natural sounds as well as 36 modulation-matched synthetics. Of these 36 natural sounds, there were 8 speech stimuli and 10 music stimuli, two of which contained singing (these stimuli were designed prior to the discovery of a song-selective component and so were not designed to examine song selectivity). Using the electrode weights from the 165 natural sounds experiment, we inferred the response of the same components to the new sound set, thus providing an independent validation of their selectivity. We plot the response timecourse of each component to natural and modulation-matched sounds separately for speech, sung music, instrumental music, and all other non-speech and non-music sounds (**Fig 3B**), as well as the time-averaged response for each pair of natural and modulation-matched sounds (**Fig 3C**).

For all category-selective components, we observed a much stronger response to natural sounds compared to synthetic sounds with matched modulation statistics. The speech-selective components (C1, C15) replicated their selectivity for natural speech (with an intermediate response to sung music) and produced weak responses to modulation-matched speech (p < 0.01 via a sign test across sounds comparing natural and modulation-matched speech). The music-selective component (C10) replicated its selectivity for natural music and responded only weakly to modulation-matched music (p < 0.01 via a sign test comparing natural and modulation-matched music). Finally, the song-selective component (C15) responded nearly exclusively to the natural sung music with almost no response to natural speech, natural instrumental music, and the modulation-matched sung music (p < 0.01 via a sign test comparing natural and modulation-matched music with singing; because there were only 2 sung music stimuli, the response to those two stimuli was subdivided into 500 ms segments to increase the number of samples). This finding demonstrates that speech, music and song selectivity cannot be explained by standard frequency and modulation statistics.

### Components selective for standard acoustic features

We next describe 6 other reliable components whose response correlated with standard acoustic features (**Fig 4A**), demonstrating that category-selective responses account for only a portion of human cortical organization, as expected (Norman-Haignere et al., 2015). These components had weights that clustered in and around primary auditory cortex (**Fig 4B**) and had fast response latencies (71-124 ms, excluding C14 which responded to sound offset). Responses to natural and matched synthetic sounds were more similar than that found for the category-selective components (**Fig 4C**), suggesting that frequency and modulation statistics account for more of their response. Most of the response variance in these components could be explained by a strong response at sound onset or offset, the magnitude of which varied across the sound set (**Fig S11**). To test if this variation could be predicted by standard acoustic features, we correlated the strength of each component’s onset/offset response – measured as the first principal component of the sound x time matrix (**Fig S11**) – with acoustic measures of audio frequency (**Fig 4D**) and spectrotemporal modulation energy (**Fig 4E**).

**Figure 4.**
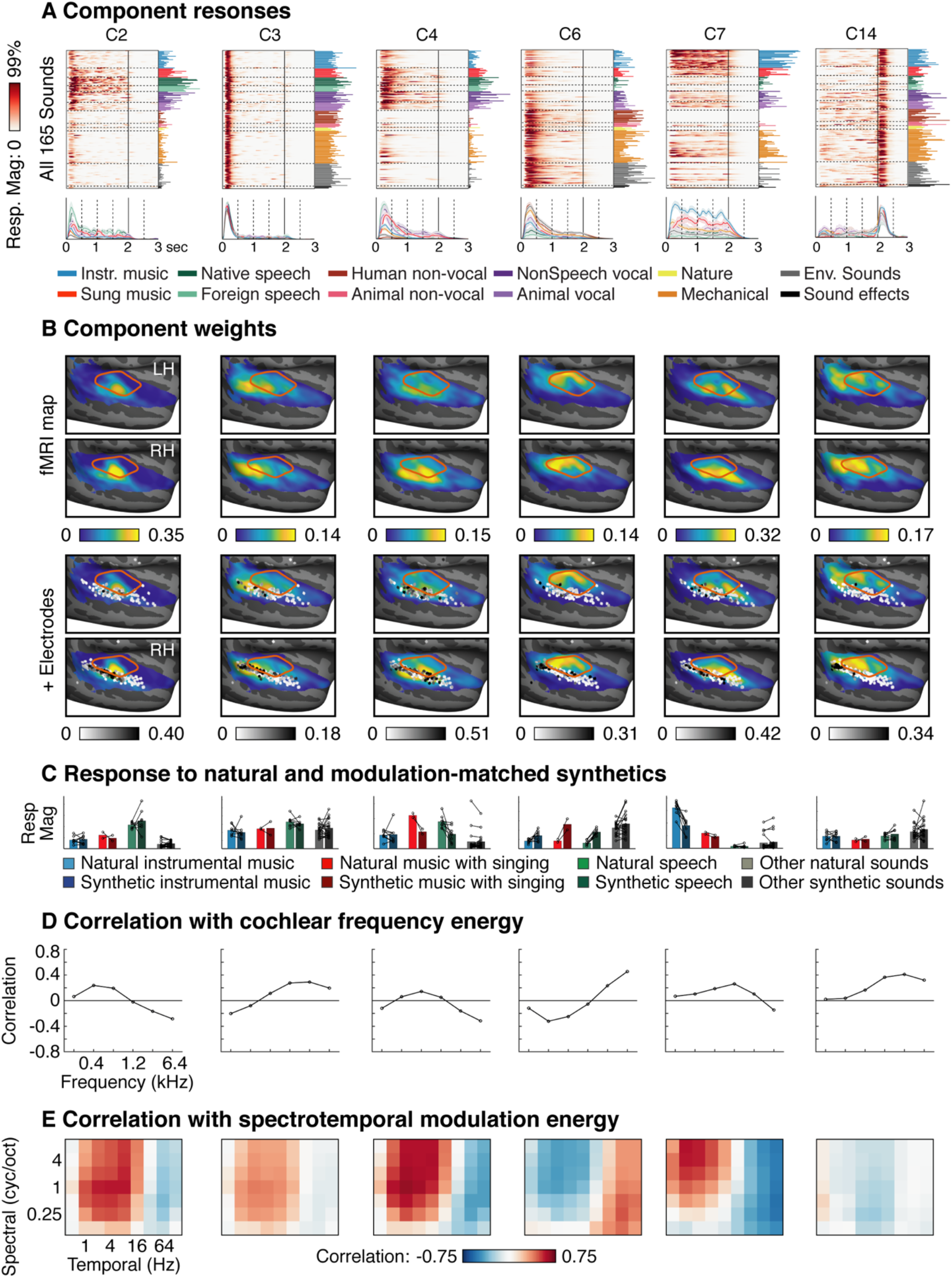
Components selective for standard acoustic features. **A-B** Responses and anatomical distribution for 6 components whose responses suggested selectivity for standard acoustic features. Same format as Figure 3A-B. **C**, Component responses to pairs of natural and modulation-matched synthetic sounds (pairs indicated by connected circles) grouped by sound category. Same format Figure 4C. **D-E**, Correlations between component responses and measures of audio frequency (panel D) and spectrotemporal modulation energy (panel E), computed from a cochleagram representation of sound. See text for details.

C3 showed a strong onset response for nearly all sounds and had weights that clustered in posterior auditory cortex, replicating prior findings of strong onset responses in this region (Hamilton et al., 2018). C14 responded strongly at sound offset, and also had weights clustered in posterior auditory cortex, which to our knowledge is the first demonstration of an anatomically organized offset response in human auditory cortex. C2 & C6 partially reflected tonotopic organization (Humphries et al., 2010; Costa et al., 2011; Moerel et al., 2012; Baumann et al., 2013): the response variation of these components correlated with measures of low and high-frequency energy and their weights clustered in low- and high-frequency regions of primary auditory cortex. These two components also showed inverse modulation tuning with C2 responding preferentially to slow temporal modulations and C6 responding preferentially to fast modulations, consistent with prior studies (Santoro et al., 2014; Norman-Haignere et al., 2015). C7 replicated prior findings of “pitch” or “tone” selectivity (Patterson et al., 2002; Penagos et al., 2004; Norman-Haignere et al., 2013): the response of this component correlated with spectrotemporal modulation energy at fine spectral scales and slow temporal rates, which is characteristic of tonal sounds, and the anatomy of this component partially overlapped the low-frequency region of primary auditory cortex and extended into more anterior non-primary regions, consistent with prior findings. Finally, C4 responded preferentially to sounds with vocal content (speech, human and animal vocalizations, and music with singing), and had weights that extended from primary regions into the middle STG, replicating prior findings of voice selectivity (Belin et al., 2000). C4 was also the only component that showed significant right-lateralization (p < 0.05 after Bonferroni correction for multiple components), consistent with prior work showing right-lateralized voice selectivity (Belin et al., 2000) (**Fig S8**). Both C4 & C7 showed modest selectivity for natural vs. synthetic sounds (for instrumental music in C7 and for speech and music with singing in C4) and had anatomical weights that straddled the border of primary/non-primary auditory cortex, suggesting a mixture of lower- and higher-order selectivity.

### Single-electrode analyses

We next tested if we could observe evidence for speech, music and song selectivity in individual electrodes without statistical assumptions or modeling. Based on prior studies, we expected that speech selectivity would be prominent in individual electrodes, but it was unclear if we would observe any selectivity for music and song in individual electrodes given that we have previously found music selectivity to be weak in individual fMRI voxels (Norman-Haignere et al., 2015; Boebinger et al., 2020).

Using a subset of data, we identified electrodes selective for speech, music or song, and then measured their response in independent data. The electrode selection stage involved three steps (all performed on the same data and distinct from that used to measure the response). First, we measured the average response across time and stimuli to all sound categories with more than five exemplars. Second, we identified a pool of electrodes with a highly selective (selectivity > 0.6) and significant (p < 0.001 via bootstrapping) response to either speech, music or song. Selectivity was measured by contrasting the maximum response across all speech and music categories (English speech, foreign, speech, sung music, instrumental music) with the maximum response across all other non-music and non-speech categories ([A-B]/A where is A is the maximum category response and B is the maximum response across all non-music/non-speech categories). Third, from this pool of music- or speech-selective electrodes, we formed three groups: those that responded significantly more to speech than all else (max[English speech, foreign speech] > max[non-speech categories except sung music]), music than all else (instrumental music > max[non-music categories]), or that exhibited super-additive selectivity for singing (sung music > max[English speech, foreign speech] + instrumental music) (p < 0.01 via bootstrapping).

We plot the response of the top electrodes most significantly responsive to each contrast (**Fig 5A**) as well as the average response across all electrodes identified using this procedure (**Fig 5B**). We measured responses to the same natural sounds used to identify the electrodes (in independent data), as well as the natural and synthetic sounds from our control experiment (**Fig 5C**). As expected, we observed a large number of speech-selective electrodes (173 electrodes across all 14 subjects). But notably, we also observed a small number of music and song-selective electrodes (11 music-selective electrodes across 4 subjects, and 7 song-selective electrodes across 3 subjects). Despite their small number, each music and song-selective electrode identified replicated their selectivity for music or speech in independent data (p < 0.05 for every electrode; p < 0.001 for responses averaged across all music and song-selective electrodes; via bootstrapping the same contrast used to select electrodes but in independent data); and modulation-matched synthetic sounds produced a much weaker responses than natural sounds from the preferred category (p < 0.01 via a sign test between responses to natural and model-matched sounds applied to the average response of speech, music, and song-selective electrodes). The subjects where we observed music and song-selective electrodes tended to be those subjects with the best coverage and the greatest number of sound-responsive electrodes, suggesting that music and song-selective electrodes would be present in most subjects if ECoG coverage were better (of the 4 subjects with the most electrodes, 3 showed music and song-selective electrodes).

**Figure 5.**
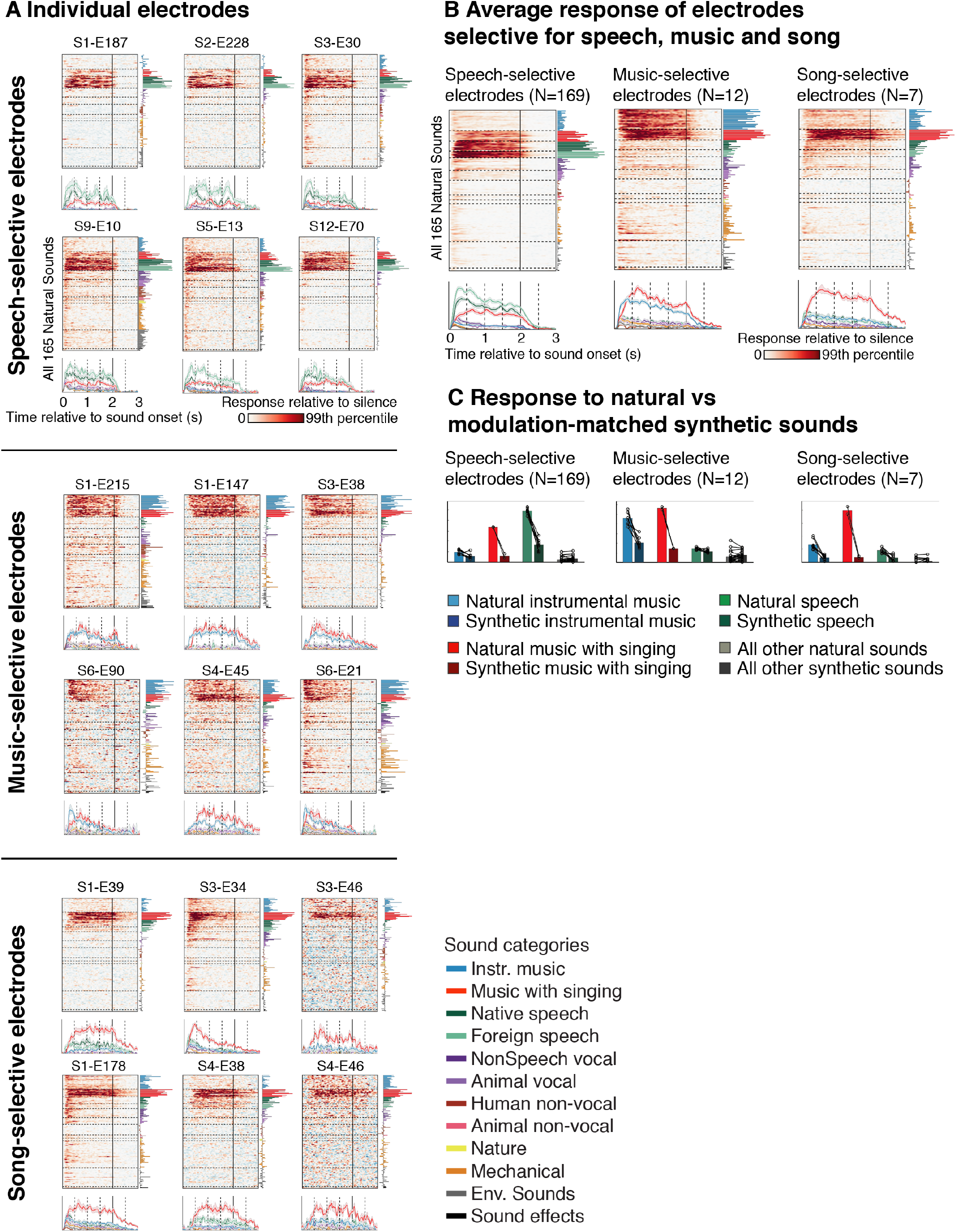
The response of individual electrodes selective for speech, music or song. We selected speech (top), music (middle), and song-selective (bottom) electrodes, and then measured their response in independent data. **A**, The top six electrodes that showed the most significant response preference for each category in the subset of data used for electrode selection. For speech-selective electrodes, the top 6 electrodes came from 2 subjects (2 from S1 and 4 from S2), and so we instead plot the top electrode from 6 different subjects to show greater diversity. Same format as in Figure 3. **B**, The average response of all electrodes identified as speech, music, or song-selective to the 165 natural sounds. **C**, The average response of speech, music, and song-selective electrodes to natural and modulation-matched synthetic sounds. Same format as Figure 4C.

The fact that we observed selectivity for singing in individual electrodes confirms that our component analysis did not infer a form of selectivity not present in the data. At the same time, song selectivity was clearly weak in individual electrodes: only a handful of electrodes showed song selectivity, and the selectivity of these electrodes was substantially weaker than the song-selective component we identified (p < 0.001 via bootstrapping the super-additive song selectivity metric), even though the component was identified using only statistical criteria without any information about the sounds tested. This observation suggests that our component method was able to isolate selectivity for song by de-mixing weak song selectivity present in individual electrodes. To more directly test this hypothesis, we re-ran our component analysis after discarding all song-selective electrodes identified in our previous analysis (note two song-selective electrodes were already discarded from our original analysis because their test-retest reliability fell below our cutoff). This analysis revealed a nearly identical song-selective component (**Fig S12**) that was again much more selective than any individual electrode (a similar music-selective component was also observed when discarding music-selective electrodes; **Fig S12**). This finding demonstrates that we can infer song selectivity using two completely non-overlapping sets of electrodes and two very different methods, one of which is blind to the properties of the stimulus. It also demonstrates the utility of component methods in identifying key response patterns that are only weakly present in individual electrodes and that one might not otherwise think to look for.

### Prediction of coarse-scale music and speech selectivity

Why were we able to observe a song-selective component that was not evident in prior fMRI studies? One natural hypothesis is that ECoG is a more precise measure of neural activity and thus allowed us to resolve finer-grained selectivity. We have already shown that song-selectivity cannot be predicted from the kinds of response components detected in our prior fMRI studies (**Fig S9**) (Norman-Haignere et al., 2015; Boebinger et al., 2020). Here, we ask the opposite: whether coarse-scale speech and music selectivity detected in our prior fMRI study can be predicted by the components identified here. To answer this question, we attempted to predict the response of the speech and music-selective components from our prior fMRI study as a weighted sum of the components identified here with ECoG, after averaging their response across time (**Fig 6**). We found that these cross-validated predictions were surprisingly accurate, accounting for 97% of the explainable response variance in both the speech and music-selective component. The fact that the fMRI components could be predicted from our ECoG components, but not vice versa (**Fig S9**), clearly demonstrates that we were able to infer finer-grained selectivity using ECoG data.

**Figure 6.**
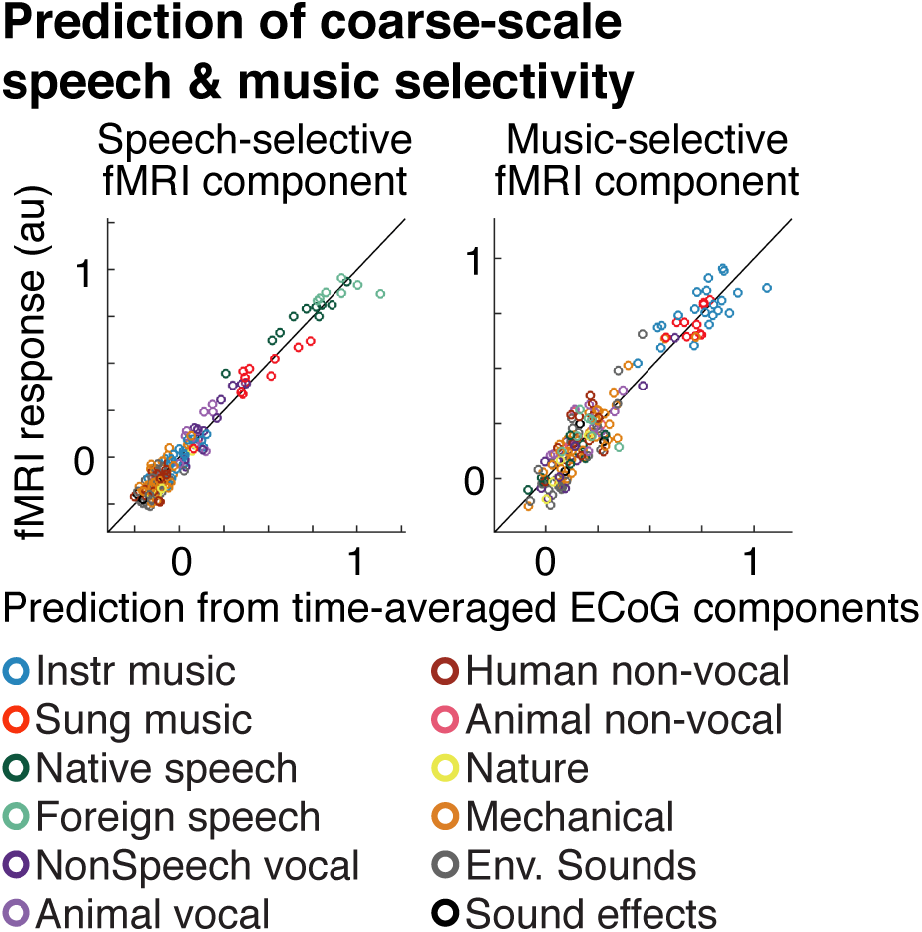
Prediction of coarse-scale speech and music-selectivity. We attempted to predict the response of the speech- and music-selective fMRI components inferred in our prior study (Norman-Haignere et al., 2015) as a weighted combination of the ECoG components identified here, after averaging their response across time. This figure plots the measured and predicted response of each component. Predictions were made using linear regression, cross-validated across the sound set (using ridge regression with five-fold, nested cross-validation).

## Discussion

Our study reveals that the human brain contains a neural population selective for song that is distinct from neural populations that respond to music and speech more generally. Song selectivity was demonstrated using a data-driven technique that was blind to the properties of sound set, a simpler regression-based method that directly searched for a linear component with binary song selectivity, and was also evident in individual electrodes. Song selectivity was co-located with music and speech-selective responses in the middle and anterior superior temporal gyrus and could not be explained by a generic acoustic model based on spectrotemporal modulation. These findings suggest that music is represented by multiple distinct neural populations, selective for different aspects of music, at least one of which responds specifically to singing.

### Song selectivity

Although song has frequently been used to explore the neural basis of music and speech perception (Merrill et al., 2012; Tierney et al., 2013; Whitehead and Armony, 2018), our findings provide the first evidence for a neural population specifically involved in the perception of song. Why might the human brain have neural populations selectively responsive to song? Vocals are pervasive in music, and typically carry the main melodic line. Thus, the brain may develop neural mechanisms specialized for representing song, simply because it is one of the first and/or most prominent components of the music that people hear. Alternatively, neural specializations for song may be partly innate, reflecting the biological importance of singing (Mehr and Krasnow, 2017).

It will be important in future work to identify the features of singing that drive song selectivity. For example, one could explore sensitivity to the types of pitch and rhythmic variation that characterize singing vs. speaking (Tierney et al., 2013). This question could be examined by using component methods to isolate selectivity for song, and then measuring the response of this component to a new sound set, as was done here to investigate selectivity for spectrotemporal modulations (**Figure 3**).

### Music selectivity

Previously, we used voxel decomposition of fMRI responses to infer a component that was substantially more selective for music than were individual voxels, which we hypothesized was due to the overlap of distinct neural populations within a voxel (Norman-Haignere et al., 2015; Boebinger et al., 2020). The present study validates these prior findings by showing the existence of music-selective neural populations using both component methods (**Fig 2&6**) and single-electrode analyses (**Fig 5**). Moreover, many of the individual electrodes that showed the strongest selectivity for music (e.g. S1-E147, S1-E215) were sampled by a high-density grid with particularly small electrodes (1 mm exposed diameter), suggesting that high spatial resolution is important for detecting music selectivity in individual electrodes.

### Speech and voice selectivity

Many prior studies have reported selectivity for speech (Mesgarani et al., 2014; Norman-Haignere et al., 2015; Overath et al., 2015) and non-speech vocalizations (e.g. crying, laughing) (Belin et al., 2000) in the superior temporal gyrus. Distinguishing responses to speech and voice has been difficult, because speech-selective responses typically show at least some response to non-speech vocalizations and vice-versa. Our component analysis revealed distinct components that responded selectively to speech (C1, C15) and non-speech vocalizations (C4), suggesting that speech and voice selectivity are partly dissociable in the human brain. Thus, as with the music selectivity, the fact that fMRI voxels reflect a mixture of speech and voice selectivity may in part reflect the blurring together of nearby neural populations.

### Onset/offset selectivity

Many of the components we observed responded substantially more strongly at the onset of sound. One of these components (C3) produced a strong onset response for nearly all sounds and had a clear posterior bias, consistent with a recent study (Hamilton et al., 2018). Another component (C14) responded selectively at sound offset, and also showed a posterior bias, suggesting that onset and offset responses are co-located to a similar region. Many other components showed an onset response whose strength was modulated by the acoustics of the stimulus (**Fig 4D,E**). Some of these responses might reflect a generic/low-level adaptation mechanism in response to a sudden increment or decrement in sound energy. Others might reflect adaptation to higher-level stimulus statistics (Kvale and Schreiner, 2004), perhaps in the service of creating a more noise-robust (Mesgarani et al., 2014) or efficient (Barlow, 1961; Fairhall et al., 2001) representation of sound by suppressing responses to features that are predictable (Heilbron and Chait, 2017).

### Component modeling: strengths, limitations and relationship to prior methods

Component modeling of both voxels with fMRI and time courses with ECoG provides a way to: (1) infer prominent response patterns; (2) suggest novel hypotheses that might not be obvious a priori; and (3) disentangle spatially overlapping responses. Our results illustrate each of these benefits. We were able to infer a small number of components that explained much of the response variation across hundreds of electrodes. We found evidence for a novel form of music selectivity (song selectivity) that we did not expect a priori. And the selectivity that we observed in the song selective component was often clearer than that evident in individual electrodes, many of which appeared to reflect a mixture of music, speech and song selectivity. Indeed, we observed a component with near-binary song selectivity even when we excluded all of the song-selective electrodes found in our dataset.

The key challenge of component modeling is that matrix approximation is inherently ill-posed, and hence, the solution depends on statistical assumptions. Most component methods rely on just one of the following three assumptions: (1) non-negativity (Lee and Seung, 1999); (2) sparsity across time or space (Olshausen and Field, 1997; Hyvarinen, 1999); or (3) temporal smoothness (Wiskott and Sejnowski, 2002; Byron et al., 2009). We showed that all of these properties are evident in auditory ECoG responses. We developed a simple model to embody these assumptions and showed that the model better predicted ECoG responses compared with baseline models. We also showed that all of our category-selective components were evident using a model that imposed only non-negativity on the responses, suggesting that our key results were robust to the particular statistical assumptions imposed (**Fig S3**). We also provided evidence for super-additive song-selectivity using a simple regression-based analysis (**Fig S10**) and by analyzing individual electrodes, neither of which depend on statistical assumptions like non-negativity or sparsity.

Our prior fMRI voxel decomposition method used statistical constraints on the high-dimensional voxel weights to infer components (Norman-Haignere et al., 2015). By contrast, ECoG grids have many fewer electrodes than voxels, but each electrode has a richly structured timecourse. We thus chose to constrain the solution with statistics of the high-dimensional response timecourses. Our method is also distinct from a number of other component models that have been applied to high-dimensional neural data. Unlike many sparse convolutional models (Bouchard et al., 2017), each component of our model was defined by a single timecourse and a single pattern of electrode weights rather than by a time-varying spatial pattern, and thus can be more easily interpreted as the response of an underlying neuronal population. Unlike clustering methods (or convex NMF (Hamilton et al., 2018)), our method can disentangle responses that overlap within individual electrodes. And unlike most tensor decomposition methods (Williams et al., 2018), our method does not require the shape of a component’s response timecourse to be identical across different stimuli, which is critical for modeling responses to sensory features that are not necessarily aligned to stimulus onset.

### Combining the strengths of fMRI and ECoG data

fMRI and ECoG data have very different strengths and weaknesses. fMRI data is coarse due to the indirect sampling of neural activity via blood flow, but it is non-invasive and can provide dense, comprehensive coverage of the entire human brain from a large number of subjects. ECoG coverage by contrast is sparse and driven by clinical demands, but has much better precision due to the direct sampling of electrophysiological activity. Our study shows one way to combine the strengths of these two methods, by inferring a canonical set of canonical response patterns with ECoG and then mapping their spatial distribution with fMRI. This approach obviously has limitations, since for example we cannot use fMRI to spatially distinguish two components with similar time-averaged responses (e.g. the speech-selective components C1 and C15). But empirically, we found a close correspondence between fMRI and ECoG maps that was primarily limited by the sparse coverage of the ECoG data (**Fig S7**). By combining the strengths of ECoG and fMRI data, we were able to detect a neural component selective for song that went undetected in our prior fMRI studies and provide an estimate of its spatial distribution in auditory cortex.

### Conclusions and future directions

By revealing a neural population selective for song, our study begins to unravel the neural code for music in the human brain, raising many questions for future research. What features of music underlie selective responses to music and song? Do these responses reflect note-level structure (e.g. pitch and timbre) (Casey et al., 2012) or the way notes are patterned to create music (e.g. melodies, harmonies and rhythms) (Schindler et al., 2013)? How can we describe the tuning of music and song-selective neural populations in computational terms, given that standard acoustic features appear insufficient (Kell et al., 2018)? And how is music and song selectivity constructed over the development of each individual, and over the history of our species (Wallin et al., 2001)? The findings and methods presented here provide a path towards answering these longstanding questions.

## Methods

### Participants

Fifteen epilepsy patients participated in our study (mean age: 37 years, age standard deviation: 14 years; 8 right-handed; 8 female). These subjects underwent temporary implantation of subdural electrode arrays at Albany Medical College to localize the epileptogenic zones and to delineate these zones from eloquent cortical areas before brain resection. All of the subjects gave informed written consent to participate in the study, which was approved by the Institutional Review Board of Albany Medical College.

### Electrode grids

Most subjects had electrodes implanted in a single hemisphere, and STG coverage was much better in one of the two hemispheres in all subjects (8 right hemisphere patients and 5 left hemisphere patients; **Fig 1B** shows the coverage of the primary hemisphere for all subjects). In most subjects, electrodes had a 2.3 mm exposed diameter with a 6 mm center-to-center spacing for temporal lobe grids (10 mm spacing for grids in frontal, parietal and occipital lobe, but electrodes from these grids typically did not show reliable sound-driven responses; electrodes were embedded in silicone; PMT Corp., Chanhassen, MN). Two subjects were implanted with a higher-density grid (1 mm exposed diameter, 3 mm center-to-center spacing). One subject was implanted with stereotactic electrodes instead of grids.

### Natural sounds

The sound set was the same as in our prior study (Norman-Haignere et al., 2015). To generate the stimulus set, we began with a set of 280 everyday sounds for which we could find a recognizable, 2-second recording. Using an online experiment (via Amazon’s Mechanical Turk), we excluded sounds that were difficult to recognize (below 80% accuracy on a ten-way multiple choice task; 55–60 participants for each sound), yielding 238 sounds. We then selected a subset of 160 sounds that were rated as most frequently heard in everyday life (in a second Mechanical Turk study; 38–40 ratings per sound). Five additional ‘‘foreign speech’’ sounds were included (‘‘German,’’ ‘‘French,’’ ‘‘Italian,’’ ‘‘Russian,’’ ‘‘Hindi’’) to distinguish responses to acoustic speech structure from responses to linguistic structure (the 160-sound set included only two foreign speech stimuli: “Spanish” and “Chinese”). In total, there were 10 English speech stimuli, 7 foreign speech stimuli, 21 instrumental music stimuli, and 11 music stimuli with singing (see *Sound category assignments*). Sounds were RMS-normalized and presented at a comfortable volume using sound isolating over-the-ear headphones (Panasonic RP-HTX7, 10 dB isolation). The sound set is freely available:

http://mcdermottlab.mit.edu/svnh/Natural-Sound/Stimuli.html

Subjects completed between three and seven runs of the experiment (S11: 3 runs, S6, S12: 4 runs, S13: 5 runs, S1: 7 runs; all other subjects: 6 runs). In each run, each natural sound was presented at least once. Between 14 and 17 of the sounds were repeated exactly back-to-back, and subjects were asked to press a button when they detected this repetition. This second instance of the sound was excluded from the analysis, because the presence of a target could otherwise bias responses in favor of the repeated stimuli. Each run used a different random ordering of stimuli. There was a 1.4-2 second gap (randomly chosen) between consecutive stimuli.

### Modulation-matched synthetic sounds

In ten of the subjects, we also measured responses to a distinct set of 36 natural sounds and 36 corresponding synthetic sounds that were individually matched to each natural sound in their spectrotemporal modulations statistics using the approach described in Norman-Haignere & McDermott (2018). The synthesis algorithm starts with an unstructured noise stimulus, and iteratively modifies the noise stimulus to match the modulation statistics of a natural sound. Modulations are measured using a standard model of auditory cortical responses in which a cochleagram is passed through a set of linear filters tuned to modulations at a particular audio frequency, temporal rate, and spectral scale (i.e. how coarse vs fine the modulations are in frequency) (Chi et al., 2005). The spectrotemporal filters were created by crossing 9 temporal rates (0.5, 1, 2, 4, 8, 16, 32, 128 Hz) with 7 spectral scales (0.125, 0.25, 0.5, 1, 2, 4, 8 cycles per octave), and replicating each filter at each audio frequency. The synthesis procedure alters the noise stimulus to match the histogram of response magnitudes across time for each filter in the model, which has the effect of matching all time-averaged statistics (such as mean and variance) of the filter responses. The stimuli and synthesis procedures were very similar to those used in Norman-Haignere & McDermott with a few minor exceptions that are noted next.

All stimuli were 4 seconds long. We used shorter stimuli than the 10-second stimuli used in Norman-Haignere & McDermott (2018) due to limitations in the recording time. Because the stimuli were shorter, we did not include the very low-rate filters (0.125 and 0.25 Hz), which were necessary for longer stimuli to prevent energy from clumping unnaturally at particular moments in the synthetic recording. We also did not include “DC filters” as in Norman-Haignere & McDermott, but instead simply matched the mean value of the cochleagram across time and frequency at each iteration of the algorithm. Norman-Haignere & McDermott describe two versions of the algorithm: one in which the histogram-matching procedure was applied to the raw filter outputs and one where the matching procedure was applied to the envelopes of the filter responses. We found that the resulting stimuli were very similar, both perceptually and in terms of the cortical response. The stimuli tested here were created by applying the histogram matching procedure to the envelopes.

The stimuli were presented in a random order with a 1.4-1.8 second gap between stimuli (for the first subject tested, a 2-2.2 second gap was used). The natural sounds were repeated to make it possible to assess the reliability of stimulus-driven responses. For all analyses, we simply averaged responses across the two repetitions. The sound set was presented across 4 runs. In one subject (S1), the entire experiment was repeated once.

### Sound category assignments

In an online experiment, Mechanical Turk participants chose the category that best described each of the 165 sounds tested, and we assigned each sound to its most frequently chosen category (30–33 participants per sound) (Norman-Haignere et al., 2015). Category assignments were highly reliable (split-half kappa = 0.93). We chose to re-assign three sounds (“cymbal crash”, “horror film sound effects”, and “drum roll”) from the “instrumental music” category to a new “sound effects” category. There were two motivations for this re-assignment: (1) these three sounds were the only sounds assigned to the music category that produced weak fMRI responses in the music-selective component we inferred in our prior study, presumably because they lack canonical types of musical structure (i.e. clear notes, melody, rhythm, harmony, key, etc.); and (2) excluding these sounds makes our song selectivity contrast (sung music – (instrumental music + speech)) more conservative as it is not biased upwards by the presence of instrumental music sounds that lack rich musical structure.

### Preprocessing

Preprocessing was implemented in MATLAB. The scripts used to implement the preprocessing steps are available here (we reference specific scripts within these directories in describing our analyses):

https://github.com/snormanhaignere/ecog-analysis-code
https://github.com/snormanhaignere/general-analysis-code

The responses from all electrodes were common-average referenced to the grand mean across all electrodes (separately for each subject). We excluded noisy electrodes from the common-average reference by detecting anomalies in the 60 Hz power (see channel_selection_from_60Hz_noise.m; a IIR resonance filter with a 3dB down bandwidth of 0.6 Hz was used to measure the RMS power). Specifically, we excluded electrodes whose 60 Hz power exceeded 5 standard deviations of the median across electrodes. Because the standard deviation is itself sensitive to outliers, we estimated the standard deviation using the central 20% of samples, which are unlikely to be influenced by outliers (by dividing the range of the central 20% of samples by that which would be expected from a Gaussian of unit variance; see zscore_using_central_samples.m). After common-average referencing, we used a notch filter to remove 60 Hz noise and its harmonics (60, 120, 180, and 240 Hz; see notch_filt.m; an IIR notch filter with a 3dB down bandwidth of 1 Hz was used to remove individual frequency components; the filter was applied forward and backward using filtfilt.m).

We computed broadband gamma power by measuring the envelope of the preprocessed signal filtered between 70 and 140 Hz (see bandpass_envelopes.m; bandpass filtering was implemented using a 6^th^ order Butterworth filter with 3dB down cutoffs of 70 and 140 Hz; the filter was applied forward and backward using filtfilt.m). The envelope was measured as the absolute value of the analytic signal after bandpassing. For the single-electrode analyses (**Fig 5**), we downsampled the envelopes to 100 Hz (from the 1200 Hz recording rate), and smoothed the timecourses with a 50 ms FWHM kernel to reduce noise and make it easier to distinguish the timecourses for different categories in the plots. For component analyses, we downsampled the envelopes to 25 Hz, because this enabled us to fit the data in the limited memory available on the GPU used to perform the optimization (no smoothing was applied since the model inferred an appropriate smoothing kernel for each component).

Occasionally, we observed visually obvious artifacts in the broadband gamma power for a small number of timepoints. These artifacts were typically localized to a small fraction of electrodes; thus, we detected artifacts separately for each electrode. For each electrode, we computed the 90^th^ percentile of its response magnitudes across all timepoints, which is by definition near the upper-end of that electrode’s response distribution, but which should also be unaffected by outliers assuming the number of outliers accounts for less than 10% of time points (which we generally found to be the case). We classified a timepoint as an outlier if it exceeded 5 times the 90^th^ percentile value for each electrode. We found this value to be relatively conservative in that only a small number of timepoints were excluded (<1% for all sound-responsive electrodes), and the vast majority of the excluded timepoints were artifactual based on visual inspection of the broadband gamma timecourses. Because there were only a small number of outlier timepoints, we replaced the outliers values with interpolated values from nearby non-outlier timepoints. We also manually excluded some or all of the runs from 11 electrodes where there were a large number of artifacts.

For each 2-second stimulus, we measured the response of each electrode during a three-second window locked to sound onset (for the 4-second natural and modulation-matched stimuli we used a 5-second window). We detected the onset of sound in each stimulus by computing the waveform power in 10 ms bins (with a 2 ms hop), and selecting the first bin in which the audio power exceeded 50 dB of the maximum power across all windows and stimuli. Following standard practice, the audio and ECoG data were synced by sending a copy of the audio signal to the same system used to record ECoG signals. This setup allowed us to measure the time delay between when the system initiated a trial and the onset of sound (which we measured using pure tones).

Responses were converted to units of percent signal change relative to silence by subtracting and then dividing the response of each electrode by the average response during the 300 ms before each stimulus.

### Session effects

For five of the thirteen subjects, runs were collected across two sessions with a gap in between (typically a day; the 7th run of S1 was collected in a third session). For the vast majority of electrodes, we found that their response properties were stable across sessions. Occasionally, we observed electrodes that substantially changed their selectivity, potentially due to small changes in the positioning of the electrodes over the cortex. To identify such changes, from each run of data, we measured the time-averaged response of each electrode to each of the 165 sounds tested. We then detected electrodes for which the test-retest correlation from runs of the same session was significantly greater than the test-retest correlation from runs of different sessions (p < 10^−5^; we used time-averaged response profiles rather than the raw timecourses, because we found them to be more reliable, and thus a better target for detecting selectivity changes across sessions; for S1 we grouped the runs from the 2^nd^ and 3^rd^ session together since there was only a single run in the 3^rd^ session). Significance was evaluated via a permutation test (Nichols and Holmes, 2002) in which we permuted the correspondence between stimuli across runs (10,000). We used this approach to build up a null distribution for our test statistic (the difference between the correlation within and across sessions), fit this null distribution with a Gaussian (so that we could estimate small p-values that would have been impossible to estimate via counting), and used the null to calculate a two-sided p-value (by measuring the tail probability that exceeded the test statistic and multiplying by 2). Seven electrodes passed our conservative significance threshold. For these electrodes, we simply treated the data from different sessions as coming from different electrodes, since they likely sampled distinct neural populations.

### Electrode selection

We selected electrodes with a reliable response to the sound set. Specifically, we measured the test-retest correlation of each electrode’s broadband gamma response timecourse across all sounds, measured in two splits of data (odd and even runs). We kept all electrodes with a test-retest correlation greater than 0.2 (electrodes with a test-retest correlation less than 0.2 were quite noisy upon inspection). Results were similar using a more liberal threshold of 0.1.

### Electrode localization

We localized electrodes in order to be able to visualize the electrode weights for each component. Electrode locations played no role in the identification of components or category-selective electrodes.

Following standard practice, we identified electrodes as bright spots on a post-operative computer tomography (CT) image. The CT was the aligned to a high-resolution, pre-operative magnetic resonance image (MRI) using a rigid-body transformation. We then projected each electrode onto the cortical surface, computed by Freesurfer from the MRI scan. This projection is error-prone because far-away points on the cortical surface can be spatially nearby due to cortical folding. As a consequence, it was not uncommon for electrodes very near STG, where sound-driven responses are common, to be projected to a spatially nearby point on middle temporal or supramarginal/inferior frontal gyrus, where sound-driven responses are much less common (**Fig S13**). To minimize gross errors, we preferentially localized sound-driven electrodes to regions where sound-driven responses are likely to occur. Specifically, using a recently collected fMRI dataset, where responses were measured to the same 165 sounds in a large cohort of 20 subjects with whole-brain coverage (our prior published study only had partial brain coverage (Norman-Haignere et al., 2015)), we calculated the fraction of subjects that showed a significant response to sound at each point in the brain (p < 10^−5^, measured using a permutation test (Norman-Haignere et al., 2016)). We treated this map as a prior and multiplied it by a likelihood map, computed separately for each electrode based on the distance of that electrode to each point on the cortical surface (using a Gaussian error distribution; 10 mm FWHM). We then assigned each electrode to the point on the cortical surface where the product of the prior and likelihood was greatest (which can be thought of as the maximum posterior probability solution). We smoothed the prior probability map so that it would only effect the localization of electrodes at a coarse level, and not bias the location of electrodes locally, and we set the minimum prior probability to be 0.05 to ensure every point had non-zero prior probability. We plot the prior map and the effect it has on localization in **Fig S13**.

### Responses statistics relevant to component modeling

Our component model approximated the response of each electrode as the weighted sum of a set of canonical response timecourses (“components”). The component timecourses are shared across all electrodes, but the weights are unique. We modeled each electrode as the weighted sum of multiple components because each electrode reflects the pooled activity of many neurons. When the electrode response timecourses are compiled into a matrix, our analysis corresponds to matrix factorization: approximating the data matrix as a product of a component response matrix and a component weight matrix.

Matrix factorization is inherently ill-posed (that is, there are many equally good approximations). Thus, we constrained our factorization by additional statistical criteria. Most component methods rely on one of three statistical assumptions: (1) non-negativity (Lee and Seung, 1999); (2) a non-Gaussian distribution of response magnitudes across time or space (Olshausen and Field, 1997; Hyvarinen, 1999); or (3) temporal smoothness of the component responses (Wiskott and Sejnowski, 2002; Byron et al., 2009). We investigated each of these statistical properties in broadband gamma responses to sound (**Fig S1**) in order to determine which statistics might be useful in designing an appropriate factorization method.

To evaluate non-negativity, we measured the percent of the total RMS power accounted for by positive vs. negative responses during the presentation of sound (measured relative to 300 ms of silence preceding the onset of each sound):

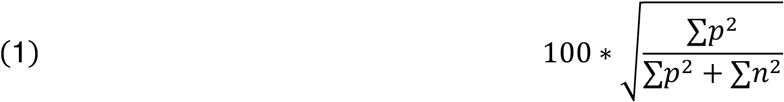

where *p* and *n* are shorthand for positive and negative values. We applied the above equation to the response of individual electrodes (summing over all timepoints for all sounds; **Fig S1A,B**), as well as to the pooled response of all sound-responsive electrodes (summing over all timepoints, sounds, and electrodes; **Fig S1D**). To minimize the effect of measurement noise, which will create negative values even if none are present (since measurement noise will not depend on the stimulus and thus noise fluctuations will be symmetric around the silent baseline), we measured the response of all electrodes in two splits of data (average across odd and even runs). We then: (1) sorted the response magnitudes of all timepoints by their magnitude in the first split; (2) measured their response in the second split; and (3) applied a median filter to the sorted response magnitudes from the second splits, thus suppressing unreliable response variation (filter size = 100 when applied to individual electrodes, filter size = 10,000 when pooling responses across all electrodes) (**Fig S1B&D** show the results of applying this procedure to individual electrodes and the pooled response of all electrodes). When equation 1 was applied to the de-noised response distributions (i.e. median filtered responses from the second split), we found that positive responses accounted for 99.97% of the RMS power across all sound-responsive electrodes. Note that sound-responsive electrodes were selected based on the reliability of their responses, not based on a greater response to sounds compared with silence, and thus our analysis is not biased by our selection criterion.

To investigate whether and how the distribution of responses might differ from a Gaussian, we measured the skewness (normalized 3^rd^ moment) and sparsity (excess kurtosis relative to a Gaussian) of the responses:

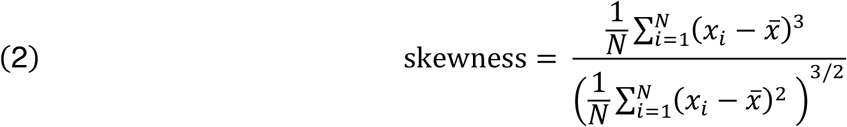

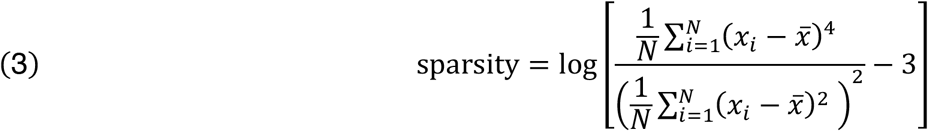

We applied the above equations to the response distribution of each electrode across all timepoints and sounds (i.e. concatenating the timecourses from all sounds into a single vector), denoised using the procedure described in the preceding paragraph. **Fig S1F** plots a histogram of these skewness and sparsity values across all electrodes. We found that all electrodes were skewed and sparse relative to a Gaussian, and relative to what would be expected given just noise in the data (p < 0.001 via sign test; see *Statistics* for details). This observation implies that the response distribution of each electrode across time/stimuli has a heavy rightward tail, with a relatively small fraction of timepoints yielding large responses for any given electrode.

We also tested the skewness and sparsity of responses across electrodes by applying equations 2 and 3 to the distribution of responses across electrodes. Specifically, we measured the averaged response of each electrode to each sound, and then for each sound, we applied equations 2 and 3 to the distribution of responses across the 271 sound-responsive electrodes. **Fig S1G** plots the histogram of these skewness and sparsity measures for all 165 sounds. We did not apply our de-noising procedure since we only had 271 electrodes which made the sorting/median-filtering procedure infeasible (in contrast, for each electrode we had 12,375 timepoints across all sounds); instead we time-averaged the response of each electrode to each sound to reduce noise. We again found that this distribution was significantly skewed and sparse relative to a Gaussian and relative to what would be expected given just noise in the data (p < 0.001 via sign test).

Finally, to investigate the temporal smoothness of auditory ECoG responses, we measured the normalized autocorrelation of each electrode’s response (**Fig S1C,E**). To prevent noise from influencing the result, we correlated responses measured in independent runs (odd and even runs). This analysis revealed substantial long-term dependencies over more than a second, and the strength of these dependencies varied substantially across electrodes. This substantial variation across electrodes demonstrates that these long-term dependencies are not a by-product of measuring broadband gamma power (in simulations, we have found that our measurement procedure can resolve power fluctuations up to ~30 Hz, assuming a 70-140 Hz carrier).

Together, the results from our analysis reveal three key properties of auditory ECoG: (1) nearly all responses are positive/excitatory relative to sound onset; (2) responses are skewed/sparse across time/stimuli and electrodes; and (3) responses are temporally smooth and the extent of this smoothness varies across electrodes. We sought to design a simple component model that captures these three essential properties. We refer to this model as the “Sparse and Smooth Component” (SSC) model.

### Component model

Each electrode is represented by its response timecourse across all sounds (*e_i_*(*t*)) (**Fig S2A**). We approximate this response timecourse as the weighted sum of K component response timecourses (***r**_k_*(*t*)):

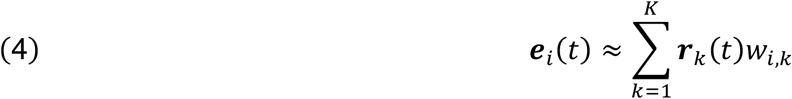

The component timecourses are shared across all electrodes, but the weights are separate for each electrode, allowing the model to approximate different response patterns. We constrain all of the component responses and weights to be positive, since we found that nearly all of the sound-driven responses were positive. To encourage the components to be both sparse and smooth, we model the response timecourse of each component as the convolution of a set of sparse activations (***α**_k_*(*t*)) with a smoothing kernel (***h**_k_*(*t*)):

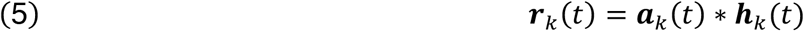

The activations effectively determine when responses occur and the smoothing kernel determines their smoothness. The activations, smoothing kernel, and electrode weights are all learned from the data. The learning algorithm proceeds by minimizing the cost function below, which has two parts: (1) a reconstruction penalty that encourages the model to be close to the data; and (2) an L1 penalty that encourages the component activations and weights to be sparse.

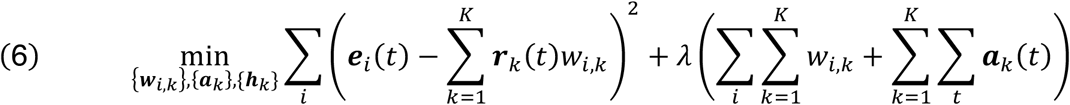

We allowed the smoothing kernel to vary across components to capture the fact that different electrodes have variable levels of smoothness. We forced the kernel to be smooth by constraining it to be unimodal (see *Constraining the Smoothing Kernel* below). The learned smoothing kernels for each component are shown in **Fig S14**. The kernels vary substantially in their extent/duration, thus capturing varying levels of smoothness across components. The model has two hyper-parameters: the number of components (*K*) and the strength of the sparsity penalty (*λ*), which we chose using cross-validation (see next section).

We implemented and optimized the model in TensorFlow, which provides efficient, general-purpose routines for optimizing models composed of common mathematical operations. We used the built-in ADAM optimizer to minimize the loss. We ran the optimizer for 10,000 iterations, decreasing the step size after each 2,000 iterations (in logarithmically spaced intervals; from 0.01 to 0.0001). Positivity of the activations and electrode weights was enforced by representing each element as the absolute value of a real-valued latent variable.

We ordered components based on their total contribution to explaining the data matrix, measured by summing the response timecourse and electrode weights for each component, and then multiplying them together:

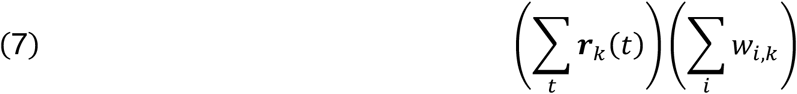

### Constraining the smoothing kernel

We investigated three potential methods for forcing the smoothing kernel to be smooth: (1) using a parametric kernel (e.g. Gamma distribution); (2) placing a smoothness penalty on the derivative of the kernel; and (3) constraining the kernel to be unimodal. We found that the optimizer had difficulty minimizing the loss when using parametric kernels (likely because the low-dimensional parameters of the kernel interacted in complex ways with the other high-dimensional parameters). We found that penalizing the derivative and constraining the kernel to be unimodal were both effective (yielding similar cross-validated prediction accuracy), but penalizing the derivative introduces a third hyper-parameter that must be chosen with cross-validation, so we chose the unimodal constraint.

We constrained the kernel to be unimodal by placing two constraints on its derivative: (1) the first N points of the derivative must be positive and the remaining points must be negative (which forces the kernel to go up and then down, but not oscillate); and (2) the sum of the derivative must equal 0 (ensuring that the kernel starts and ends at zero). The set of operations used to implement these constraints in TensorFlow is described in **Fig S15**. Many of the learned smoothing kernels were asymmetric, with a rapid rise and a slower falloff (**Fig S14**). There is nothing in the constraints that encourages asymmetry, and so this property must reflect an asymmetry in the cortical responses themselves.

### Cross-validation analyses

We used cross-validated prediction accuracy to determine the number of components and the sparsity parameter (**Fig 1E & S2B**), as well as to compare the SSC model with several baseline models (**Fig S2C**). For the purposes of cross-validation, we separated the timecourses for different sounds into cells, thus creating an electrode x sound matrix (**Fig 1E**). We then trained the model on a random subset of 80% of cells and measured the model’s prediction accuracy (squared Pearson correlation) in the left-out 20% of cells. We trained models starting from 10 different random initializations, and selected the model with the lowest error in the training data. We repeated our analyses using 5 different random splits of train and test data, averaging the test correlations across splits. For each split, we ensured an even and broad sampling of train and test stimuli using the following procedure: (1) we created a random ordering of stimuli and electrodes (2) we assigned the first 20% of sounds to be test sounds for the first electrode, the next 20% of sounds to be test sounds for electrodes 2, and so on. After using up all 165 sounds (every 8-9 electrodes), we refreshed the pool of available test sounds using a new random ordering of stimuli.

To prevent correlated noise across electrodes from influencing the results, we used non-overlapping sets of runs (odd and even runs) to compute the training and test data (i.e. training on odd runs and testing on even runs, and vice-versa; again averaging test correlations across the two splits). For a given set of hyper-parameters, we then averaged the test correlations across all electrodes to arrive at a summary measure of that model’s performance (**Fig 1E & S2B**). We noise-corrected these correlations using the test-retest correlation of each electrode’s response (see *Noise correction* below).

We considered several baseline models that did not use the convolutional decomposition of the SSC model (specifically, we constrained the smoothing kernel to be a delta function such that the component activations, ***α**_k_*(*t*), equaled the component responses, ***r**_k_*(*t*)). We tested four baseline models: (1) we removed the sparseness and smoothness constraints entirely but maintained the non-negativity constraint (i.e. non-negative matrix factorization / NMF); (2) we imposed sparsity but not smoothness via an L1 penalty the component responses and weights (3) we imposed smoothness but not sparsity via an L2 smoothness penalty on the derivative of the component responses (the first-order difference of adjacent time-points); and (4) we applied both the L1 sparsity and L2 smoothness constraint. To prevent the number of hyper-parameters from biasing the results, for each electrode, we selected the hyper-parameters that led to the best performance across electrodes from other subjects (**Fig S2C**). We used grid-search over the following range of hyper-parameters: *K* (number of components) = [5,10,15,20,25,30], *λ* (sparsity) = [0,0.033,0.1,0.33,1,3.3], *ω* (smoothness) = [0,0.033,0.1,0.33] (we verified that the best-performing models were not on the boundary of these values, except in cases where the best-performing model had a parameter value of 0). We found that all of the baseline models performed worse than the SSC model (p < 0.001 via bootstrapping across subjects, see *Statistics*; including the model with both an L1 sparsity and L2 smoothness penalty, which had more hyper-parameters). This result shows that our convolutional decomposition is an effective way of capturing both the smoothness and sparsity of auditory broadband gamma responses, and is more effective than simply imposing sparsity and smoothing penalties directly on the component responses.

### Assessing component reliability

We assessed component reliability by comparing components across models with different statistical assumptions (**Fig S3**), varying the number of components (**Fig S4**), testing whether the electrode weights were concentrated in individual subjects (**Fig S5**), and testing whether the components were robust to random re-initializations of the algorithm. We focused on a set of 10 components that were robust across all of these metrics.

We first compared components inferred by our SSC model and a simpler model that only imposed non-negativity on the responses (using a 15-component model in both cases). We greedily matched components across models by correlating their electrode weights (results were the same using response profiles). The matching process started by matching the pair of components with the highest correlation, removing those two components, and then repeated the process until no more components were left. For the 10 reliable components, the correlation of response profiles and weights for corresponding components was always higher than 0.84, and the responses were qualitatively very similar (**Fig S3**). Four components (C3, C5, C9, C13) were excluded from the main figures because they had lower response profile or weight correlations. These components are shown in **Fig S6**.

To evaluate whether the components were robust across the number of model components, we tested if the same components were present in a 20-component model (**Fig S4**). We found that most of the components present in the 15-component model were also present in a 20-component model, demonstrating that the model primarily added new components without altering existing components. For the 10 reliable components, the response profile and weight correlations were always higher than 0.88. Reducing the number of components from 15 inevitably eliminated/merged some of the components from the 15-component model (which had the best cross-validated prediction accuracy), though we note that a song-selective component still evident in a 10-component model.

For a 15-component model, we found that most components had weights that were broadly distributed across many electrodes and subjects. As we increased the number of components in the model, we found that components began to weigh heavily on a small number of electrodes often from a single subject, which may explain why cross-validation prediction accuracy dropped for higher numbers of components (**Fig 1E**). In the 15-component model, we found one component that that weighed quite heavily on a single subject (C12 weighed mainly on electrodes from S3; see **Fig S5**). The response of this component is also plotted in **Fig S6**.

As with any sparse component model, our cost function is not convex, and the optimization algorithm could potentially arrive at local optima, leading to unstable results across different random initializations of the algorithm. To address this issue, we ran the analysis many times (1,000 times), using different random initializations (activations and electrode weights were initialized with random samples from a truncated normal distribution; see **Fig S15** for the structure and initialization of the smoothing kernels). Components that are stable should be consistently present for all solutions with low cost, which we quantified by correlating the component response profiles for the solution with the lowest cost with those for the 99 next-best solutions (using the “Hungarian algorithm” to determine the correspondence between components from different solutions (Kuhn, 1955)). As a measure of stability, we computed the median correlation value for each component across the 99 next-best solutions. For all of the components, even those that were less reliable by other metrics, the median correlation was greater than 0.93, indicating that the algorithm was able to find a stable local minimum.

### fMRI weight maps

We took advantage of a large dataset of fMRI responses to the same 165 sounds in order to get a second and more reliable estimate of the anatomical weight map for our ECoG components (**Figure 2B&4B**), thus combining the broader and denser sampling available in fMRI with the more precise functional components derived from ECoG. The dataset consisted of responses from 30 subjects collected across two studies (Norman-Haignere et al., 2015; Boebinger et al., 2020). The subjects had a wide range of musical expertise from almost none to many years of training. We have found that fMRI responses to natural sounds, including selective responses to music, are very similar in subjects with and without musical training (Boebinger et al., 2020). We thus pooled data across all 30 subjects unless otherwise noted. We limited our analyses to sound-responsive voxels, as determined in our prior study(Norman-Haignere et al., 2015).

The fMRI weights were computed by approximating each voxel’s response as the weighted sum of the time-averaged ECoG components, measured using linear regression. Specifically, we multiplied the fMRI response profile of each voxel by the pseudoinverse of the time-averaged ECoG component response matrix. The time-averaged responses of the speech-selective components (C1&C15) were highly correlated (r=0.91) which would have made the pseudoinverse unstable, and thus we excluded the other speech-selective component from the pseudoinverse when calculating the projection matrix for each speech-selective component. We then averaged the weights across subjects to arrive at our group maps.

To evaluate the correspondence between the weight maps for the ECoG components derived from the ECoG versus fMRI data, we correlated weight maps for corresponding components and compared this correlation to that for mismatching components, as well as to the split-half reliability across subjects of the fMRI and ECoG maps alone (**Figure S7**). Because ECoG coverage varies from patient to patient, we carefully split the patients into two groups with a similar number of electrodes that were evenly distributed between the left and right hemisphere when evaluating split-half reliability (for fMRI analyses we simply used a random split with an even number of subjects from each study). Specifically, we considered all possible splits of the 15 ECoG patients into two groups. For each split, we computed the absolute difference between the total number of electrodes in each group as well as the absolute difference between the total number of electrodes between the left and right hemisphere of each group. We then selected the split where the sum of these two difference scores was minimal. For the optimal split, the first group had 78 right hemisphere electrodes and 58 left hemisphere electrodes across 5 patients. The second group had 68 right hemisphere electrodes and 68 left hemisphere electrodes across 10 patients (obviously with fewer electrodes per patient).

In order to compare the fMRI and ECoG maps, we resampled both to a common anatomical grid on the cortical surface (1.5 mm x 1.5 mm spacing). Because ECoG coverage varies, some grid positions are much better sampled than others. To account for this, we weighted each grid position by a measure of the total number of electrodes near to it when computing correlations. We did this for all correlations including the fMRI split-half correlations, where this is not strictly necessary, so that the results would be comparable. The weighted Pearson correlation can be computed by simply replacing the standard covariance, variance and means statistics with their weighted counterparts:

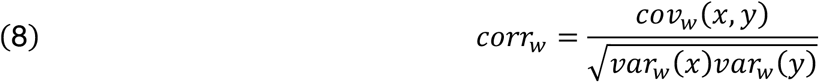

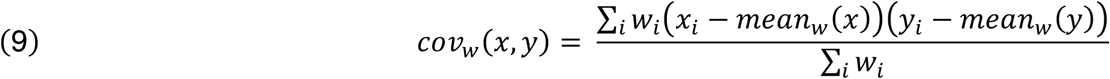

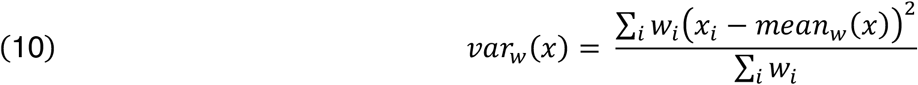

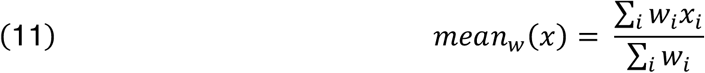

The grid weights (*w_i_*) were computed using an interpolation-like scheme, where we assigned a weight to each pair of electrodes and grid positions based on the distance between them, using a 5 mm FWHM Gaussian kernel (fMRI data were also smoothed with a 5 mm FWHM kernel). We then summed up the weights for each grid position across all electrodes. A high grid weight indicates there were many nearby electrodes, while a low weight indicates there were no nearby electrodes. We excluded electrodes with a very small weight from the analysis (w<0.1).

We bootstrapped across the fMRI subjects to get error bars for all correlations involving fMRI data (**Figure S7**). It was not possible to bootstrap the ECoG subjects, because we needed to carefully split the subjects into two groups with a similar number of electrodes.

### Component responses to modulation-matched sounds

The components were inferred using responses to just the 165 natural sounds from the main experiment. But since a subset of ten subjects were tested in both experiments, we could estimate the response of these same components to the natural and synthetic sounds from our control experiment. Specifically, we fixed the component electrode weights to the values inferred from the responses in our main experiment, and learned a new set of component response timecourses that best approximated the measured responses in the modulation-matching experiment. Since the electrode weights are known, this analysis is no longer ill-posed, and we thus removed all of the additional sparsity and smoothness constraints and simply estimated a set of non-negative response profiles that minimized the squared reconstruction error (we left the non-negativity constraint because we found that nearly all of the measured responses were non-negative).

### Acoustic correlations

We evaluated acoustic selectivity for the components that showed weak category selectivity and more similar responses to natural and modulation-matched sounds (**Fig 4**). Most of the response variance of these components could be explained by the first principal component of the sound x time response matrix (**Fig S11**), which for many of the components reflected a strong response at sound onset or offset, the magnitude of which varied across the sound set. We correlated the stimulus weights of the first PC (i.e. the magnitude of the response variation across sounds) with the same measures of frequency and spectrotemporal modulation energy used in our prior study (Norman-Haignere et al., 2015). The frequency measures were computed by summing energy in a cochleagram representation of sound across both time and frequency (within 6 coarse bands with center frequencies shown in **Fig 4D**), yielding one number per sound and frequency band. The modulation measures were computed using a standard bank of spectrotemporal modulation filters applied to a cochleagram representation of sound (Chi et al., 2005; Norman-Haignere and McDermott, 2018). This analysis yielded a 4D tensor (time x audio frequency x temporal modulation x spectral modulation) which we summed across time and audio frequency. We correlated these measures with the stimulus weights of the first PC. We partialled out the contribution of the frequency measures from the modulation measures to ensure they could not explain any modulation correlations.

### Predicting ECoG components from fMRI and vice versa

We tested if we could predict the response of the ECoG song-selective component from our previously inferred fMRI components (**Fig S9**), and whether we could predict the response of the speech and music-selective fMRI components from our ECoG components (**Fig 6**). For these analyses, we averaged electrode responses across time. Predictions were made using ridge regression with five-fold cross-validation across the sound set. The regularization parameter was chosen using cross-validation within each training fold. The folds were chosen to include a roughly equal number of sounds from each category.

We used two independent measures of each fMRI and ECoG component to noise-correct the predictions and get a measure of explainable variance (see *Noise correction* below). The two independent measurements were computed by first getting three independent measurements of each voxel or electrode response (from different repetitions). We then used one measurement to estimate the weights for each component (again using ridge regression), and then projected the data from the other two measurements onto these weights.

### Estimating a linear projection with binary song selectivity

We used a simple regression analysis to test if there was a linear projection of the electrode responses with nonlinear song selectivity (**Fig S10**). Specifically, we used ridge regression to try and learn a weighted sum of the electrode responses that yielded a binary response with 1s for all vocal music and 0s for all other sounds. We used time-averaged electrode responses to learn the weights since we only needed one number per sound to learn the mapping. We then multiplied the full timecourses of the electrodes by the learned weights in order to be able to compare the component inferred by this analysis with the component inferred by our decomposition method. We used cross-validation across the sound set to prevent statistical circularity (nested 5-fold cross-validation with the regularization parameter selected within the train set). We found that the component inferred by this analysis was very similar to the component inferred by our decomposition method, even in more detailed aspects that were not explicitly optimized for (i.e. the temporal pattern within and across vocal music stimuli was similar between the two components) (**Fig S10**).

### Single electrode analyses

To identify electrodes selective for music, speech and song, we defined a number of contrasts based on the average response to different categories (the contrasts are described in the Results). We then divided each contrast by the maximum response across all categories to compute a measure of selectivity, or we bootstrapped the contrast to determine if it was significantly greater than zero (see *Statistics* below). In all cases, we used independent data to identify electrodes and measure their response. Specifically, we used two runs (first and last) to select electrodes and the remaining runs to evaluate their response.

### Statistics

The significance of all category contrasts was evaluated using bootstrapping (Efron, 1982). Specifically, we sampled sounds from each category with replacement (100,000 times), averaged responses across the sampled sounds for each category, and then recomputed the contrast of interest (all of the contrasts tested are specified in the Results). We then counted the fraction of samples that fell below zero and multiplied by 2 to compute a two-sided p-value. For p-values smaller than 0.001, counting becomes unreliable, and so we instead fit the distribution of bootstrapped samples with a Gaussian and measured the tail probability that fell below zero (and multiplied by 2 to compute a two-sided p-value). For the component analyses, we corrected for multiple comparisons by multiplying these p-values by the number of components (corresponding to Bonferroni correction).

We compared the song-selective component (Component 12) with the average response of all song-selective electrodes by counting the fraction of bootstrapped samples where the component showed greater super-additive selectivity for singing (sung music > max(English speech, foreign speech) + instrumental music). We found that across all 100,000 bootstrapped samples, the component always showed greater selectivity.

We also used bootstrapping to compute error bars for the category timecourses (e.g. **Fig 2A**). We only plot categories for which there were more than 5 exemplars.

We used fMRI to test for laterality effects because we had a large number of subjects with complete, bilateral coverage, unlike ECoG where each patient had sparse coverage from a single hemisphere (**Fig S8**). For each component, we computed the average weight of the top 100 voxels from the left and right hemisphere, corresponding to about 10% of sound responsive voxels (1040 voxels in the right hemisphere, 991 sound-responsive voxels in the left hemisphere). We focused on the top 100 voxels because component weights were generally concentrated in a focal anatomical region. Results were robust to the specific number of voxels selected. We then subtracted the average weight for the left and right hemisphere and bootstrapped this difference score by sampling subjects with replacement (100,000 samples). We computed a p-value by counting the fraction of samples falling below or above zero (whichever was smaller) and multiplying by 2. We then Bonferroni-corrected by simply multiplying the p-value by the number of components.

We also used bootstrapping across subjects to place error bars on model prediction scores. Specifically, we (1) sampled subjects with replacement (10,000 times); (2) averaged the test correlation values across the electrodes from the sampled subjects; and (3) noise-corrected the correlation using the test-retest reliability of the sampled electrodes. We tested whether the SSC model outperformed our baseline models by counting the fraction of bootstrapped samples where the average test predictions were lower than each baseline model and multiplying by 2 to arrive at a two-sided p-value. When plotting the test predictions for different models (**Fig S2C**), we used “within-subject” error bars (Loftus and Masson, 1994), computed by subtracting off the mean of each bootstrapped sample across all models before measuring the central 68% of the sampling distribution. We multiplied the central 68% interval by the correction factor shown below to account for a downward bias in the standard error induced by mean-subtraction (Loftus and Masson, 1994):

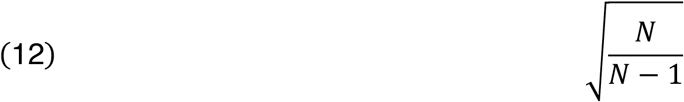

We used a sign test to evaluate whether the response to natural sounds was consistently greater than responses to corresponding modulation-matched sounds. A sign test is natural choice, because the natural and modulation-matched sounds are organized as pairs (**Fig 3A**). For components selective for speech / music (song selective components described in the next paragraph), we compared the time-averaged response to natural speech / music with the corresponding modulation-matched controls (there were eight speech stimuli, eight instrumental music stimuli and two music stimuli with singing). We performed the same analysis on the average response of speech and music-selective electrodes (**Fig 5C**). For both components and electrodes, the response to natural sounds of the preferred category was always greater than the response to modulation-matched sound, and thus significant with a sign test (p < 0.01).

Although there were only two music stimuli with singing in the modulation-matching experiment, the stimuli were relatively long (4 seconds). We thus subdivided the response to each stimulus into seven 500 ms segments (discarding the first 500 ms to account for the build-up in the response), and measured the average response to each segment. For both the song-selective component and the average response of song-selective electrodes, we found that for all fourteen 500-ms segments (7 segments across 2 stimuli), the response to natural sung music was higher than the response to the modulation-matched controls, and thus is significant with a sign test (p < 0.001).

To determine whether the electrode responses were significantly more skewed and sparse than would be expected given noise (i.e. to evaluate the significance of the skewness/sparsity measures described in *Response statistics relevant to component modeling*), we computed skewness/sparsity using two data quantities: (1) the residual error after subtracting the response to even and odd runs; and (2) the summed response across even and odd runs. The properties of the noise should be the same for these two quantities, but the second quantity will also contain the reliable stimulus-driven component of the response. Thus, if the second quantity is more skewed/sparse than the first quantity, then the stimulus-driven response must be more skewed/sparse than the noise. To assess skewness/sparsity across time/stimuli, we measured the skewness and sparsity (equations 2 and 3) separately for each electrode using the residual error and summed response (pooling responses across all timepoints and stimuli). In every subject, we found that the average skewness/sparsity of the summed responses was greater than the skewness/sparsity of the residual error, and thus significant with a sign test (p < 0.001). We used the same approach to evaluate the skewness/sparsity of responses across electrodes, measured separately for each sound. Using a sign test across sounds, we found both the skewness and sparsity of the summed response to be significantly greater than that for the residual error (p < 0.001).

### Noise correction

We used standard noise correction procedures to provide a ceiling on our measured correlations and provide an estimate of explainable variance. In general, the upper bound on the correlation between two variables is given by the geometric mean of the reliability of the variables (Schoppe et al., 2016; Kell et al., 2018):

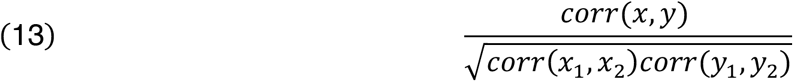

where *x*_1_ and *x*_2_ are two independent measures of the same variable. The numerator was typically computed by averaging the cross-variable correlations for two independent measurements:

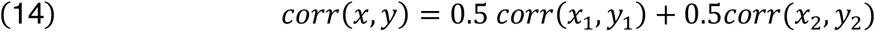

When comparing ECoG and fMRI weight maps we used all of the available data for computing the ECoG-fMRI correlations (**Fig S7**), and we therefore spearman-brown corrected the test-retest correlations in the denominator to account for the greater amount of data when calculating the noise ceiling (since we used all of the data to compare ECoG and fMRI maps but only half of the data to calculate reliability) (Spearman, 1910):

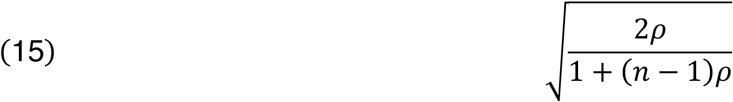

where *ρ* is the uncorrected split-half correlation.

For simplicity, when measuring the response variance explained by the components (**Fig 1E, S2B-C**), we only corrected for the reliability of the individual electrodes and not the components, since the components were much more reliable than the individual electrodes.

We used the squared and noise-corrected Pearson correlation as a measure of explainable variance.

## Acknowledgements

This work was supported by the National Institutes of Health (EY13455 to N.G.K., P41-EB018783 to G.S., P50-MH109429 to G.S., R01-EB026439 to G.S., U24-NS109103 to G.S., U01-NS108916 to G.S., and R25-HD088157 to G.S.), the U.S. Army Research Office (W911NF-15-1-0440 to G.S.), the National Science Foundation (Grant BCS-1634050 to J.H.M.), the NSF Science and Technology Center for Brains, Minds, and Machines (CCF-1231216), Fondazione Neurone (Grant to G.S.), and the Howard Hughes Medical Institute (LSRF Postdoctoral Fellowship to S.N.H.).

---

**Figure S1.**
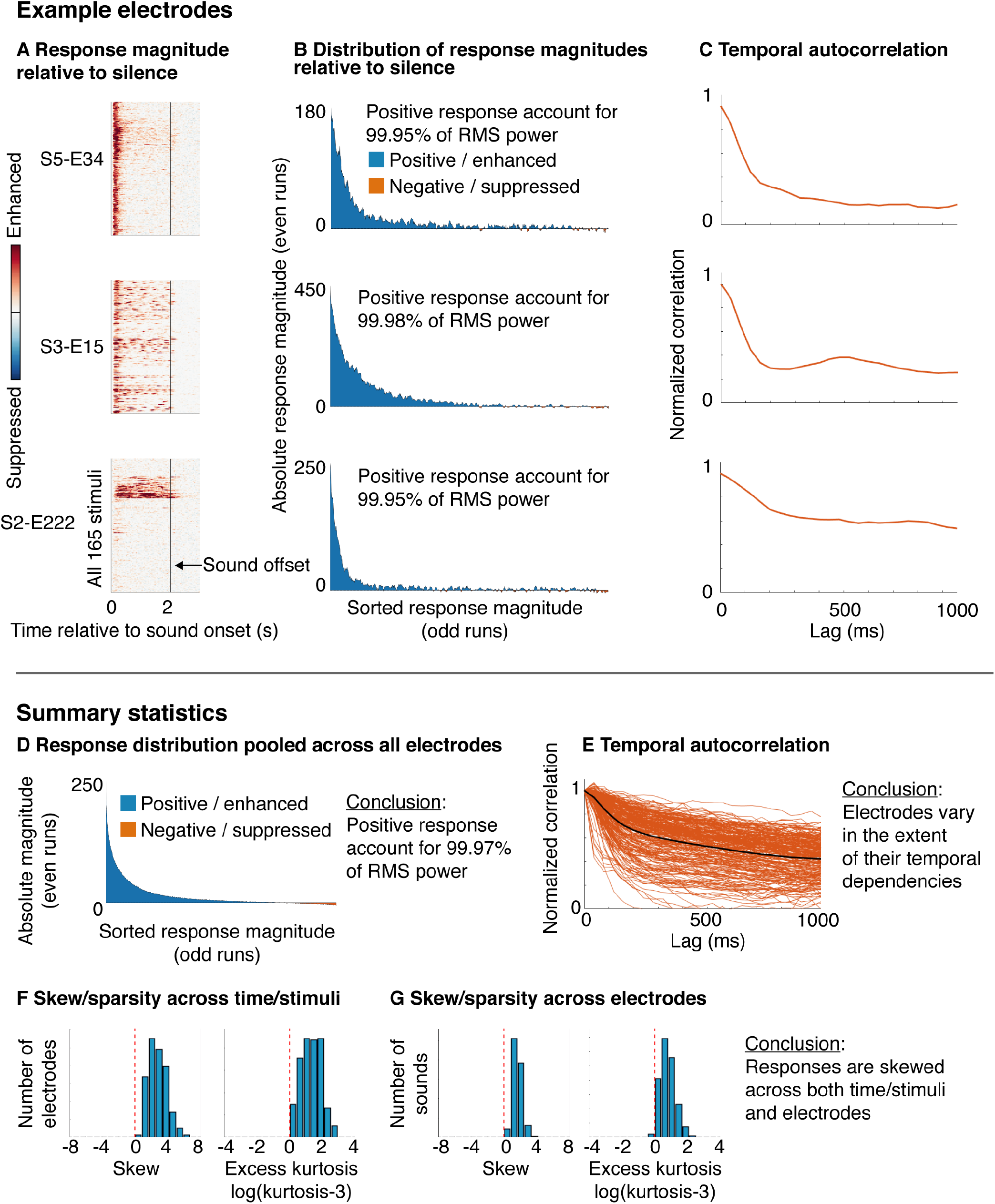
Response statistics relevant to component modeling. **A-C**, Response statistics from three example electrodes with distinct selectivities, but a shared set of statistical properties (positivity, sparsity/skew, and temporal smoothness). **A**, Broadband gamma power response of each electrode to all 165 sounds as a raster. Responses are measured relative to the response during silence (300 milliseconds preceding sound onset). Positive values (red) indicate an enhanced response to sound, and negative responses indicate a suppressed response (blue). The color scales shows values from 0 to the 99^th^ percentile of the response magnitude distribution for each electrode. **B**, Distribution of response magnitudes, measured in a cross-validated fashion to reduce effects of noise: using data from the odd runs, we sorted all of the bins of the raster on the left based on their magnitude (pooling across all timepoints and stimuli). The response of each bin was then measured using the even runs, and then smoothed using a median filter to suppress noise. Positive responses accounted for >99% of the RMS response power in all three electrodes. All three electrodes show a skewed and sparse distribution of response magnitudes (quantified in panel F, below) because negative responses were practically non-existent (yielding an asymmetric, rightward-skewed distribution) and strong positive responses were present for only a small fraction of bins (yielding a sparse distribution). **C**, The normalized autocorrelation (normalized by the correlation at zero lag) of each electrode’s response measured in a cross-validated fashion by correlating the response in odd and even runs at different lags. **D-G**, Summary statistics across all sound-responsive electrodes. **D**, Distribution of response magnitude pooled across all electrodes, sounds and timepoints (measured in a cross-validated fashion, as described above). Positive responses accounted for >99% of the RMS power. **E**, Normalized autocorrelation of all sound-responsive electrodes. The extent of temporal dependencies varied substantially across electrodes. **F**, We measured the skew (3^rd^ moment) and sparsity (excess kurtosis) of each electrode’s response using its distribution of response magnitudes across all timepoints/stimuli (i.e. using the distributions shown in panel B). This figure plots a histogram of the skew and sparsity values across all electrodes. We subtracted the measured kurtosis from that which would be expected from a Gaussian (which has a kurtosis of 3). All electrodes were skewed and sparse relative to a Gaussian. **G**, For each sound, we measured the skew and sparsity of responses across electrodes, after averaging the response of each electrode to each sound. This figure plots a histogram of the skew and sparsity values across all sounds.

**Figure S2.**
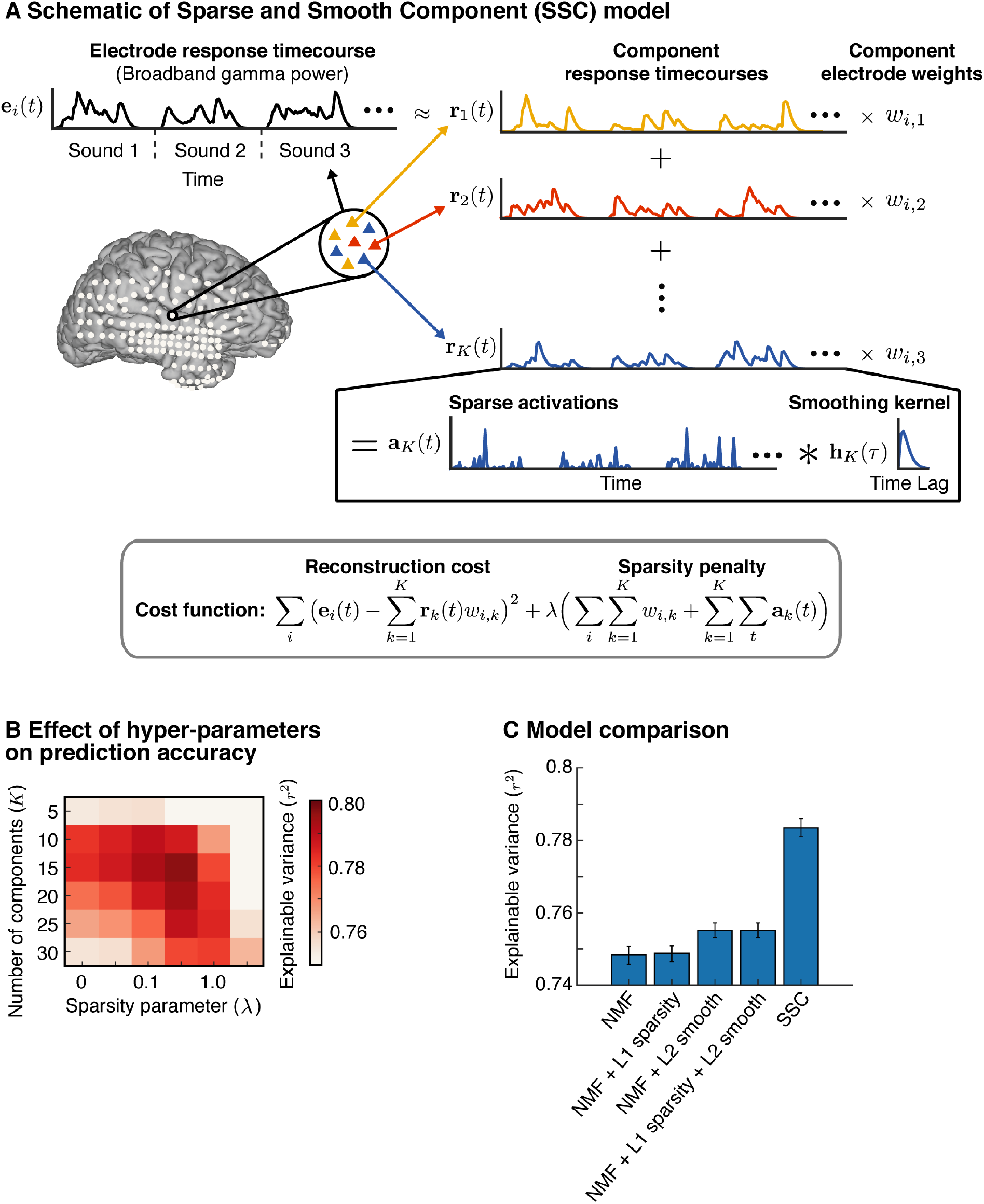
Component model and its evaluation via cross-validation. **A**, Schematic of the “sparse and smooth” component model, which was motivated by the statistical properties shown in Figure S1. Each electrode was represented by its response timecourse (broadband gamma) across all sounds (measured relative to silence). This timecourse was modeled as the weighted sum of multiple component timecourses to capture the fact that each electrode is influenced by many neurons and thus might reflect multiple underlying neuronal populations. The component response timecourses were the same across electrodes, but the weights varied to account for different response patterns. Both the component responses and weights were constrained to be positive. To encourage the component response patterns to be sparse and skewed, we modeled each component as the convolution of a set of sparse activations with a smoothing kernel. The activations, weights and smoothing kernel were all learned by minimizing a cost function with two terms: (1) a reconstruction penalty encouraging the components to closely approximate the data; and (2) a sparsity penalty encouraging the activations and weights to be sparse. The smoothing kernel was learned separately for each component to account for variable levels of smoothness in the responses across electrodes. **B**, Average squared correlation between the measured and model-predicted response in test data as a function of the number of components and sparsity penalty (the correlation has been noise-corrected; Figure 2B shows results for the best sparsity parameter (Λ = 0.33)). **C**, Comparison of the prediction accuracy (average correlation in test data) of the SSC model with several baseline models that did not rely on the convolutional decomposition used by the SSC model: (1) non-negative matrix factorization (NMF) where the components and weights were constrained only to be positive; (2) NMF with a sparsity penalty applied directly to the responses and weights; (3) NMF with a L2 smoothness penalty applied to the derivative (first-order difference) of the component responses; and (4) NMF with both an L1 sparsity and L2 smoothness penalty. Data from independent subjects was used to select the hyper-parameters for each model and evaluate prediction accuracy. Error bars show the median and central 68 percent of the sampling distribution measured via bootstrapping across subjects.

**Figure S3.**
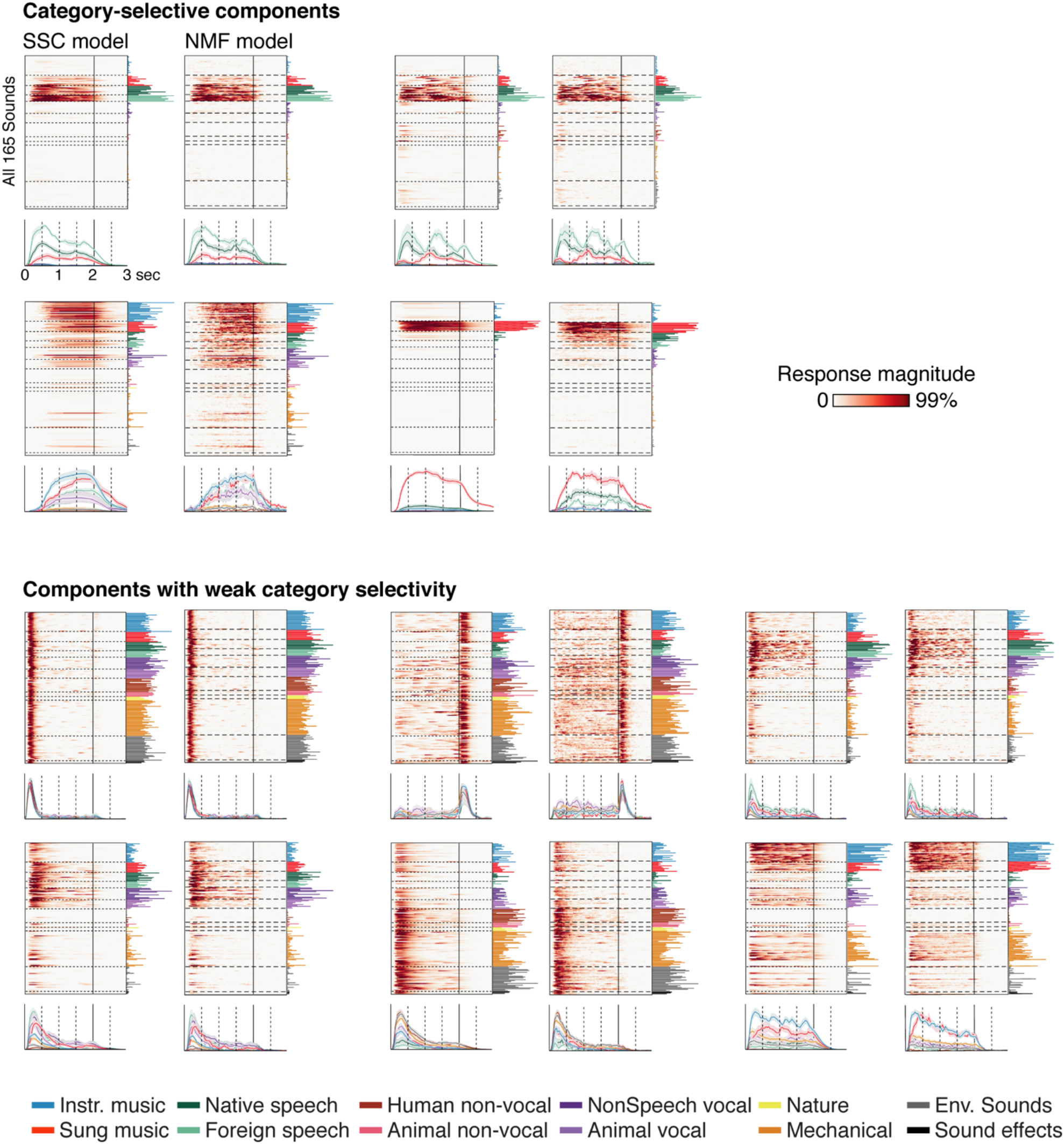
Consistency across models. The 10 reliable components estimated from our SSC component model were also evident in a model that just imposed non-negativity on the responses and electrode weights (non-negative matrix factorization or NMF). To illustrate this fact, we show corresponding components from the two models side-by-side (estimated from a 15-component model in both cases). To mirror the main text, the components are grouped into those with strong and weak category selectivity. Format is the same Figure 2A and 4A.

**Figure S4.**
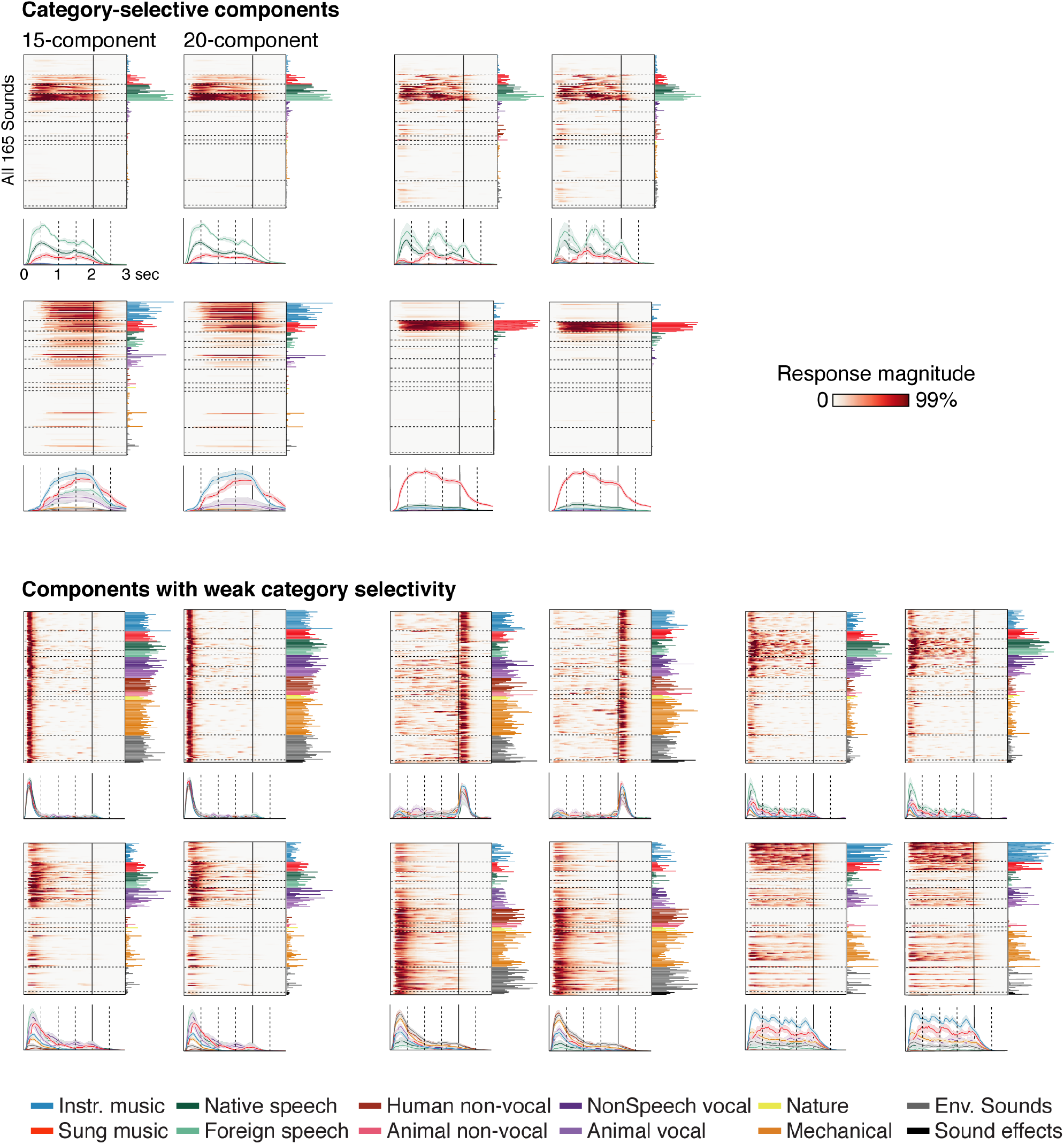
Consistency across the number of components. The 10 reliable components were clearly present in both 15- and 20-component models. To illustrate this fact, we show corresponding components from 15- and 20-component models side-by-side. To mirror the main text, the components are grouped into those with strong and weak category selectivity. Format is the same Figure 2A and 4A.

**Figure S5.**
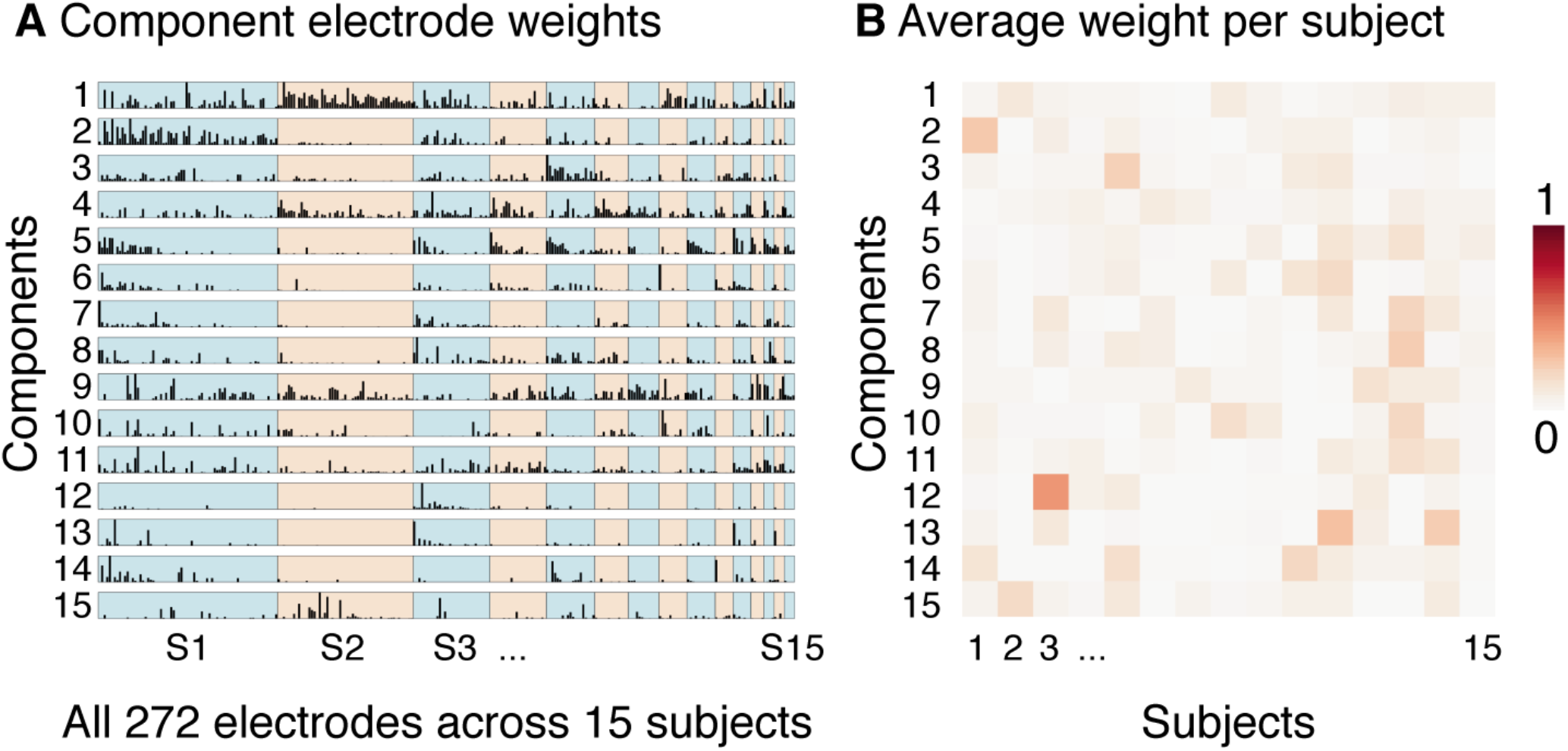
Distribution of components weights across electrodes/subjects. **A**, The distribution of weights across all electrodes for each component. Electrodes from different subjects have been grouped together, and the subjects have been ordered based on their total number of electrodes. **B**, The average weight of each component in each subject, normalized so that the weights across subjects sum to 1. As a consequence, large values indicate that a component primarily explained responses from a single subject. Most components had weights that were distributed across a large number of electrodes from several different subjects. Component 13 weighted strongly on a single subject (S3) and was not included in the figures in the main text for this reason (Figure S6 shows the less reliable components).

**Figure S6.**
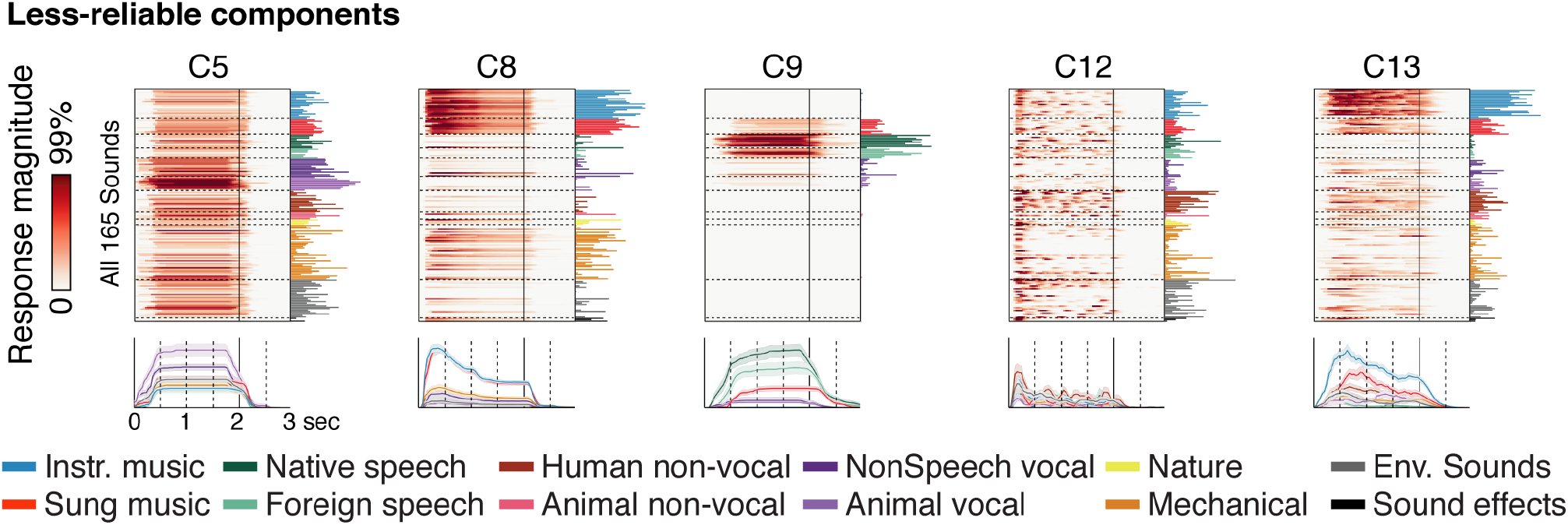
Response of less-reliable components. This figure shows the response of components that were less reliable than those plotted in the main text (same format as Fig 2A). Components C5, C8, C9 and C12 differed substantially from those evident in a simpler NMF model. C12 weighted heavily on electrodes from a single subject (Fig S6).

**Figure S7.**
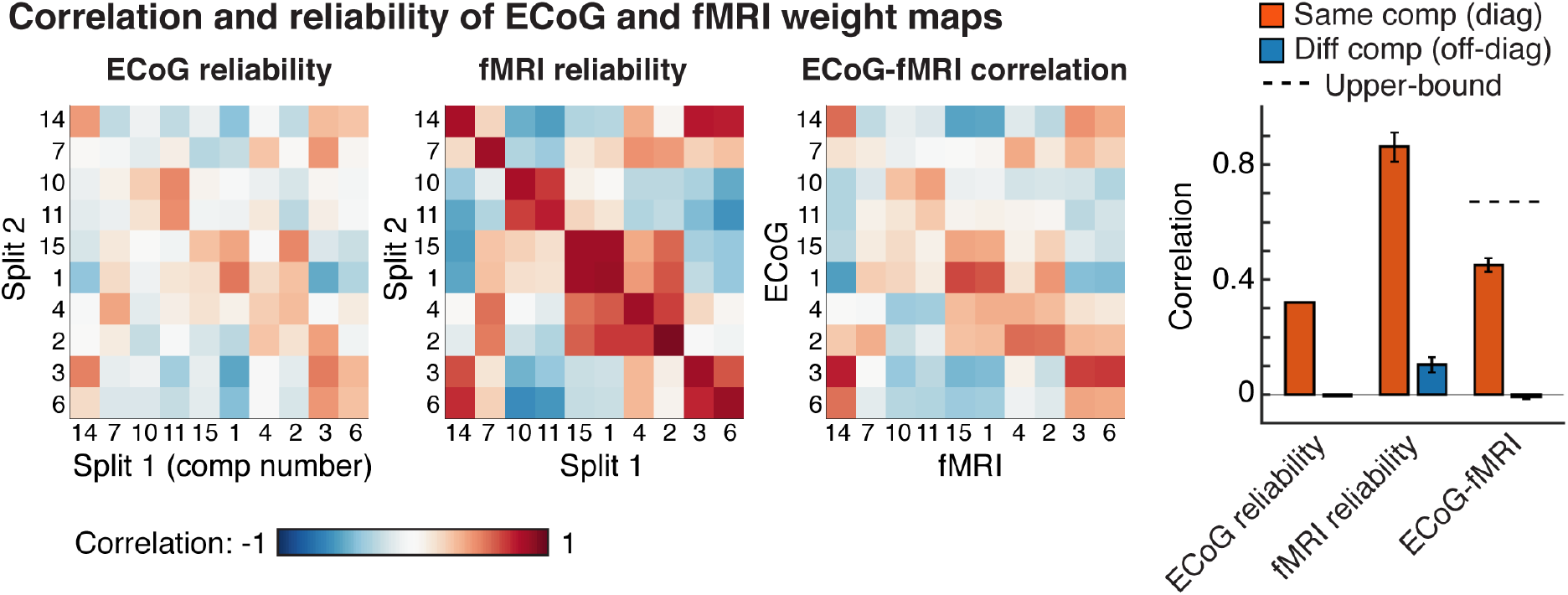
Comparing anatomical maps across fMRI and ECoG. For each inferred ECoG component, we computed an anatomical weight map using either ECoG or fMRI (Figure 2B & Figure 4B). This figure shows the split-half correlation across subjects for the ECoG (left matrix) and fMRI maps (middle matrix) as well as the correlation between ECoG and fMRI maps (right matrix). The diagonal contains the correlation across splits for corresponding components. The rightmost panels plots the average correlation for corresponding (diagonal of the matrix) and non-corresponding components (off-diagonal). If components have reliably distinct weights then the correlation for corresponding components should be higher than for non-corresponding components. Components have been ordered by the similarity of their response profile since components with more similar response profiles also tend to have more similar anatomical distributions in the brain (using a hierarchical clustering algorithm: MATLAB’s “linkage” function with correlation distance). Error bars show the central 68% of the sampling distribution computed by bootstrapping across fMRI subjects (bootstrapping across ECoG subjects was not feasible). fMRI maps were much more reliable than ECoG maps due to superior coverage, and the correlation between fMRI and ECoG maps was slightly higher than the split-half reliability of the ECoG data itself. The dashed line shows an estimate of the maximum possible correlation between ECoG and fMRI maps given the reliability of the two measures.

**Figure S8.**
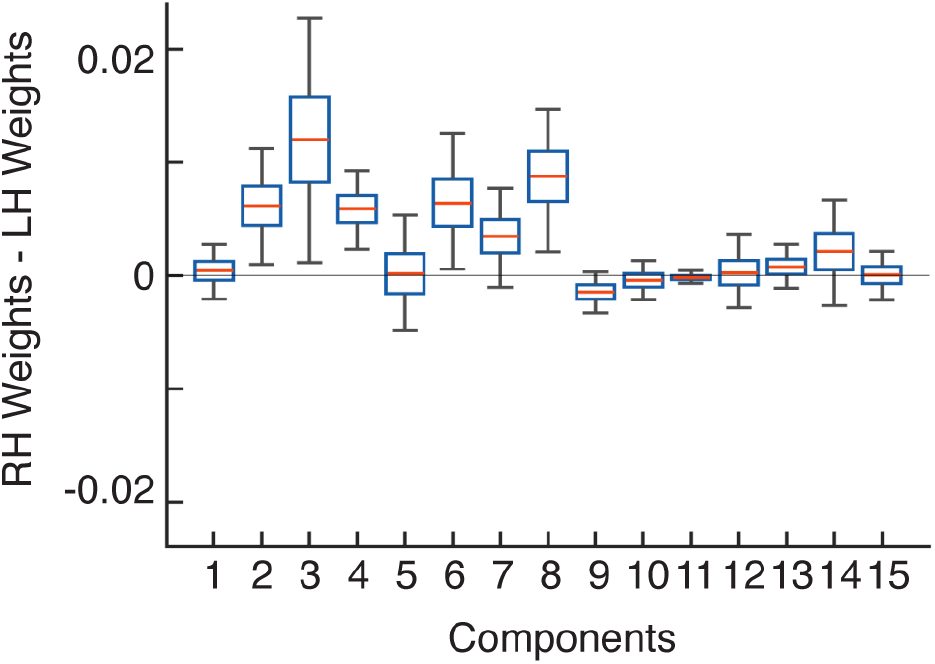
Laterality of component electrode weights. We used the fMRI weight maps, where we had good, bilateral coverage from 30 subjects, to test for differences in laterality. This figure plots the average weight difference between the left and right hemisphere. Boxes and whiskers show the central 50% and 95% of the sampling distribution, respectively, estimated via bootstrapping across subjects.

**Figure S9.**
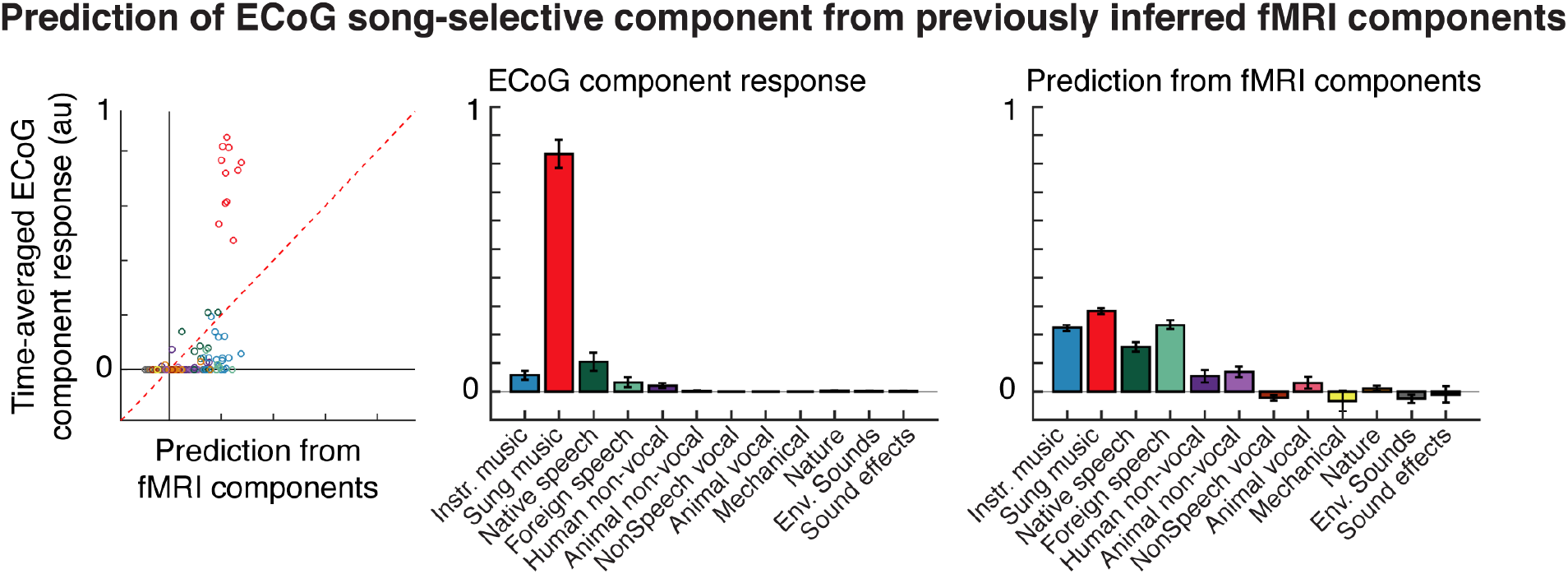
Song-selectivity cannot be predicted from fMRI components. We previously inferred a set of fMRI components that showed selectivity for speech and music, individually. To test if the song-selective component (C11) could be explained as a weighted sum of speech and music selectivity (or any other fMRI component), we regressed the fMRI components against the time-averaged response of the song-selective component (using ridge regression with five-fold, nested cross-validation across the sound set). The left panel shows a scatter of the predictions and the right panel shows the original (middle panel) and predicted (right panel) component response, averaged across categories.

**Figure S10.**
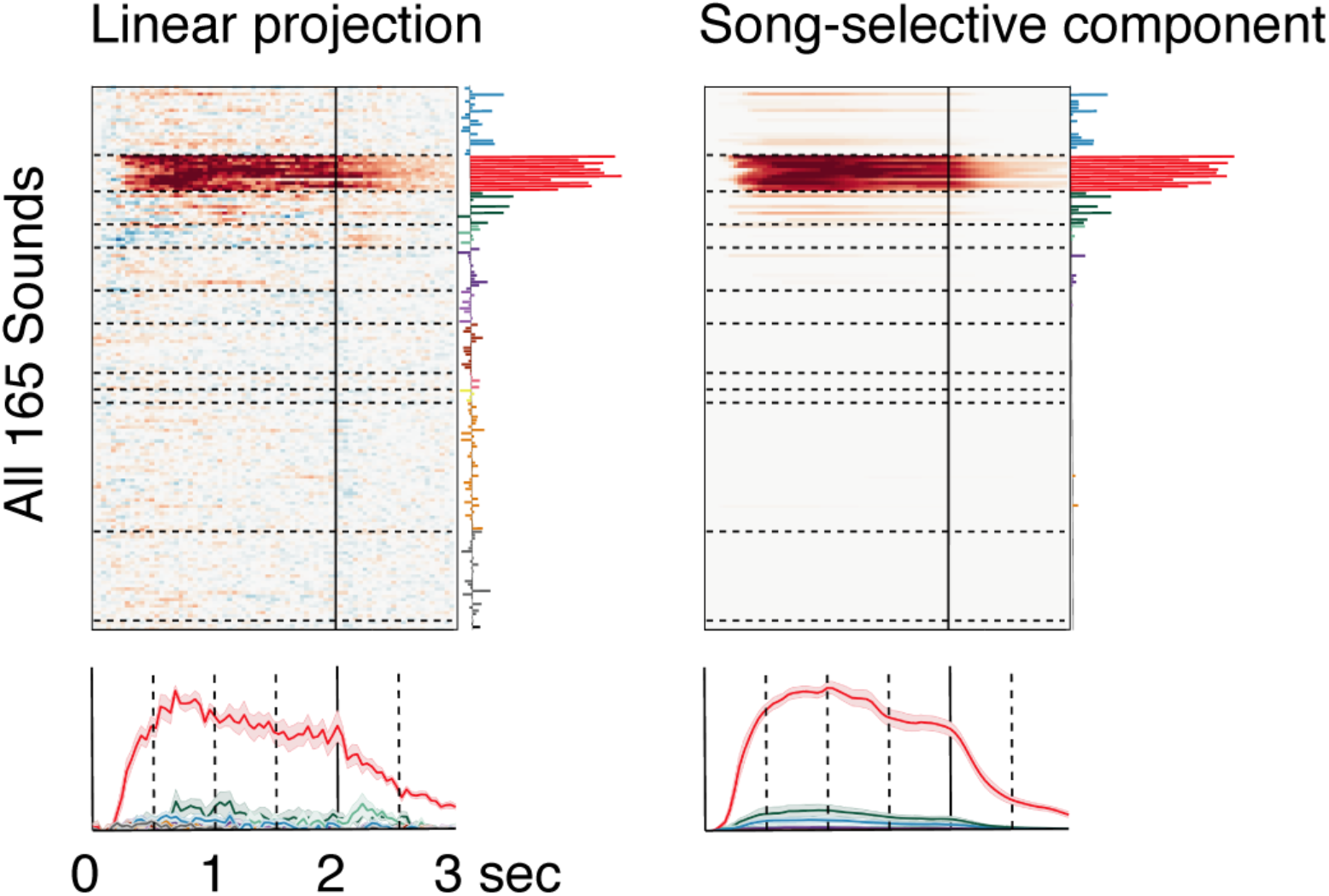
Song-selectivity computed via linear projection. The left panel plots the response of a component, inferred by finding a weighted sum of the electrodes that best approximated a binary song-selective response. Format is the same as Fig 2A. Cross-validation across sounds was used to ensure that song selectivity was genuine: weights were estimated using a subset of the sounds and responses in the left-out sounds were then projected onto these weights (using ridge regression with nested 5-fold cross-validation). The inferred component was very similar to the component inferred by our decomposition method (right panel). Because the regression-inferred component is by definition a linear function of the electrodes, its nonlinear selectivity for song cannot be explained as a linear function of speech and music selectivity present in the electrode responses.

**Figure S11.**
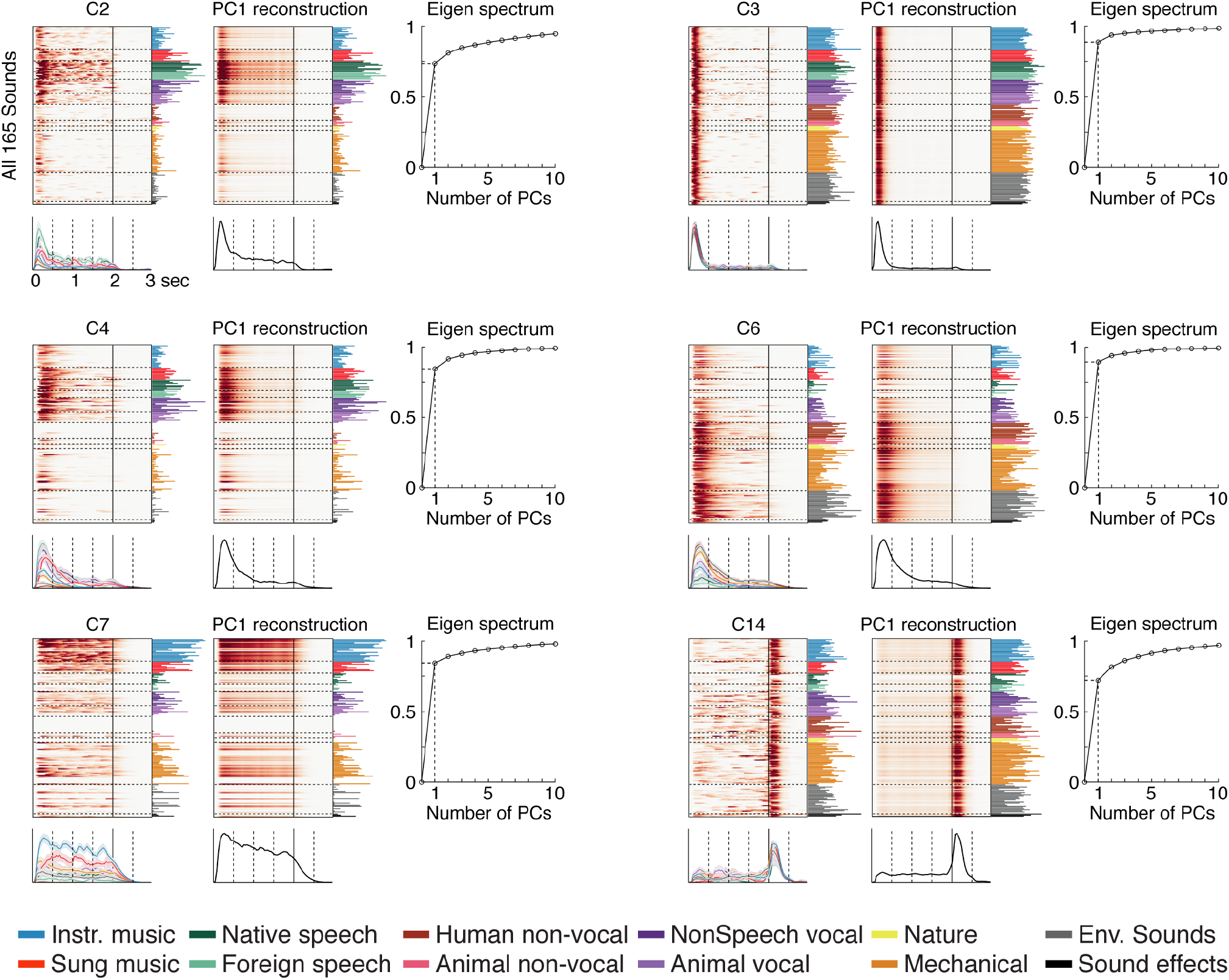
Using the first principal component to capture variation in onset/offset responses. The components with weak category selectivity (**Fig 4**) showed a strong response at the onset or offset of sound, the magnitude of which varied across the sound set. The first principal component (PC) captured much of this variation. For each component, we show the original response matrix (same as in Figure 4A) as well as the reconstruction using the first PC. Below the reconstruction, we show the timecourse of the first PC and to the right, we plot the stimulus weights for the first PC. To the right, we plot the cumulative eigen spectrum, which illustrates that the first PC accounted for much of the response variance. For our acoustic correlation analyses (Figure 4D&E), we correlated the stimulus weights of the first PC with measures of frequency and spectrotemporal modulation.

**Figure S12.**
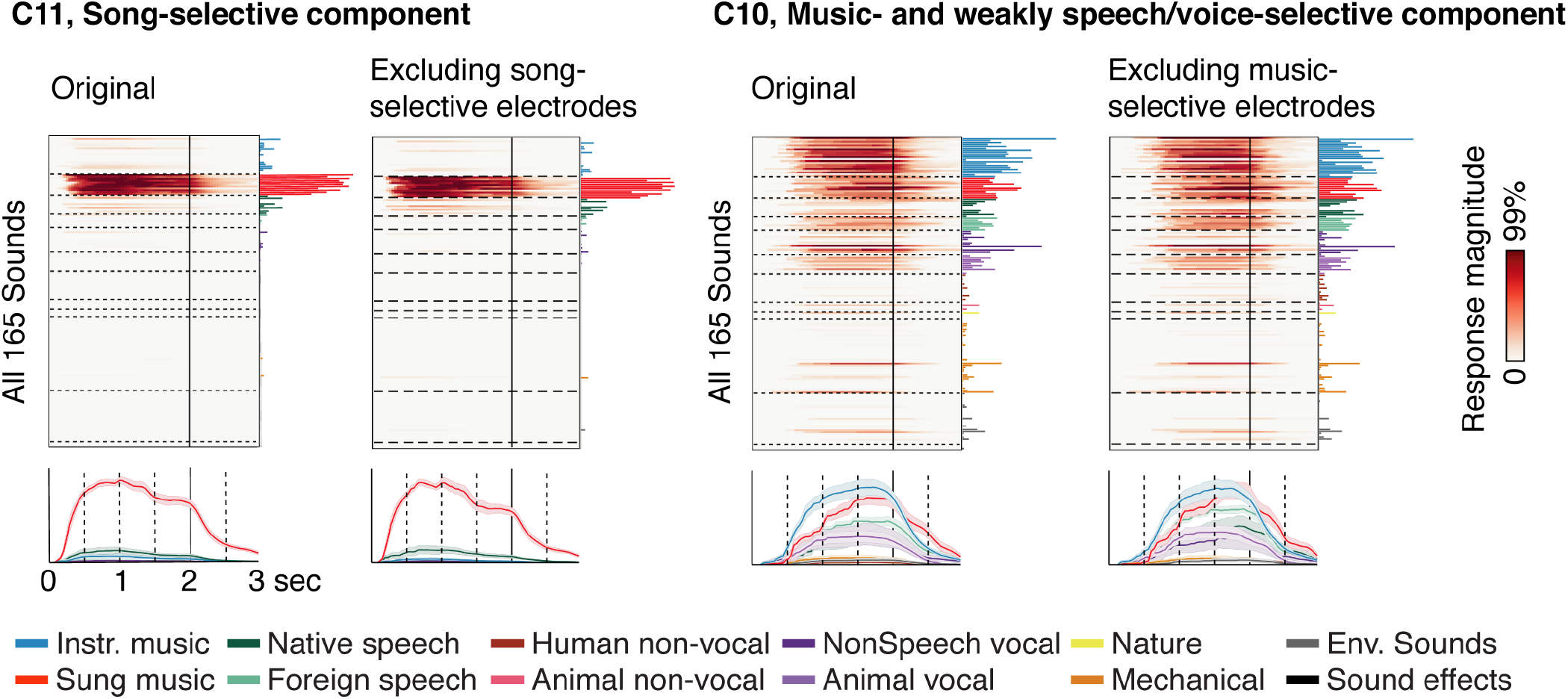
Excluding song and music-selective electrodes. This figure plots song and music-selective components inferred after throwing out all song and music-selective electrodes, respectively (same format as Figure 2A). The original components are shown side-by-side for comparison. The inferred components were largely unaffected by the presence/absence of song and music-selective electrodes.

**Figure S13.**
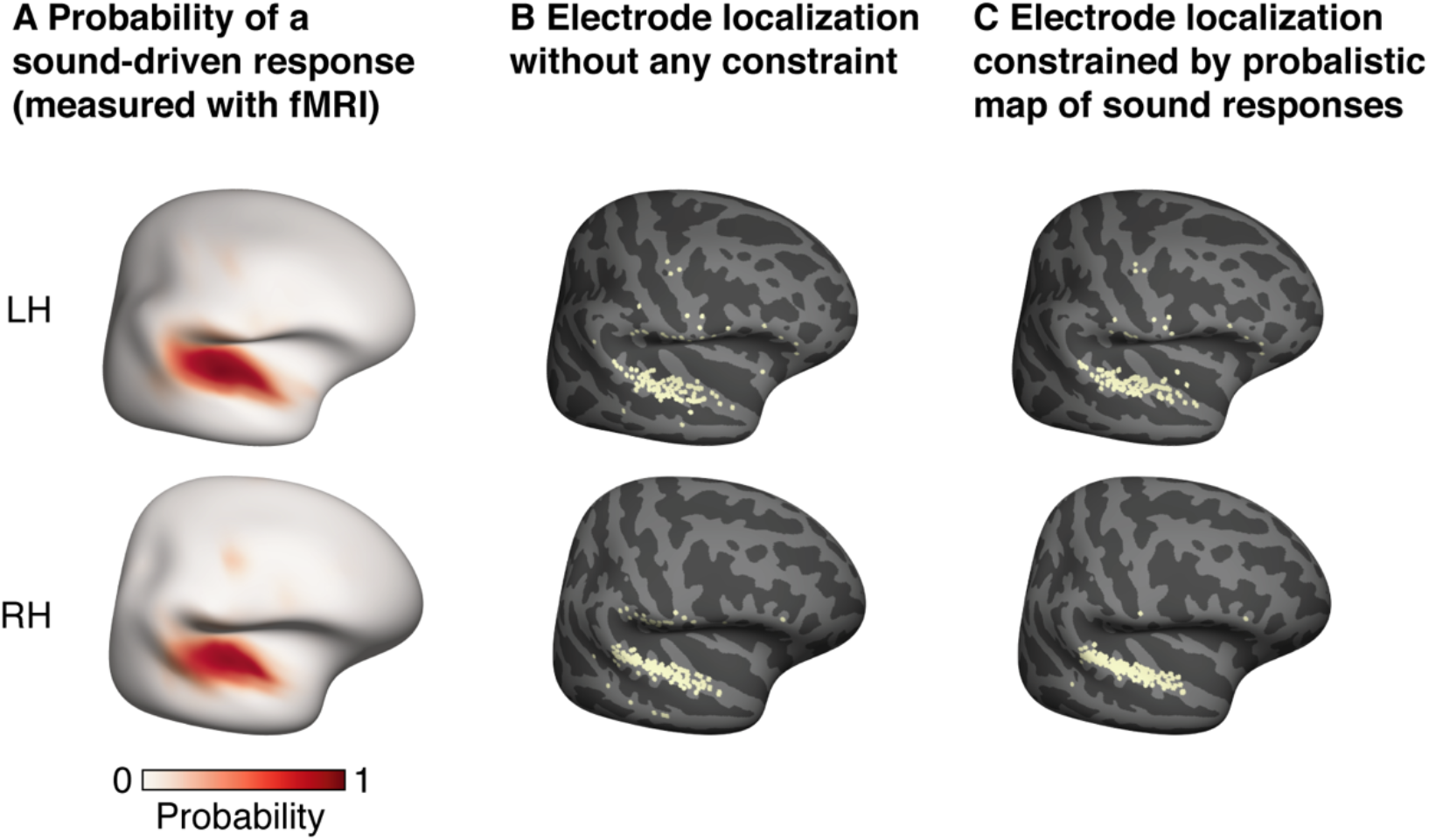
Constraining the anatomical localization of electrodes. **A**, Map showing the probability of observing a significant response to sound at each point in the brain. The map was computed using fMRI responses to the same sound set in a large cohort of 20 subjects. **B**, Electrode localization based purely on anatomical criteria. Small errors in localization likely explain why some electrodes have been localized to the middle temporal gyrus and supramarginal/inferior frontal gyrus, which abut the superior temporal gyrus where responses to sound are common. **C**, To minimize gross localization errors, we treated the probability map of sound-driven responses shown in panel A as a prior and used to it constrain the localization (see *Electrode localization* in the Methods). Our approach did not substantially affect the localization of electrodes at a fine scale, but encouraged electrodes to be mapped to the superior temporal gyrus rather than the middle temporal or supramarginal/inferior frontal gyrus.

**Figure S14.**
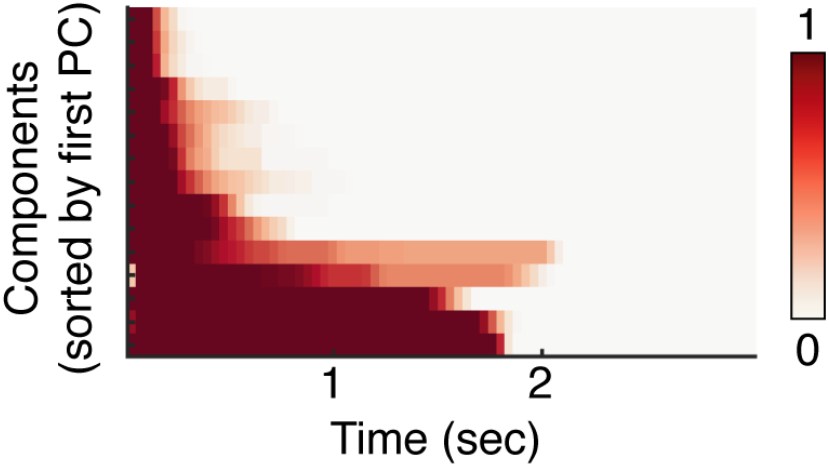
Learned smoothing kernels. This figure plots the learned smoothing kernels as a raster, with each row corresponding to a different kernel. The kernels have been sorted by the first principal component of the matrix. The kernels vary widely in their extent/duration. Many of the kernels are were also asymmetric with a fast/instantaneous rise and a slower falloff.

**Figure S15.**
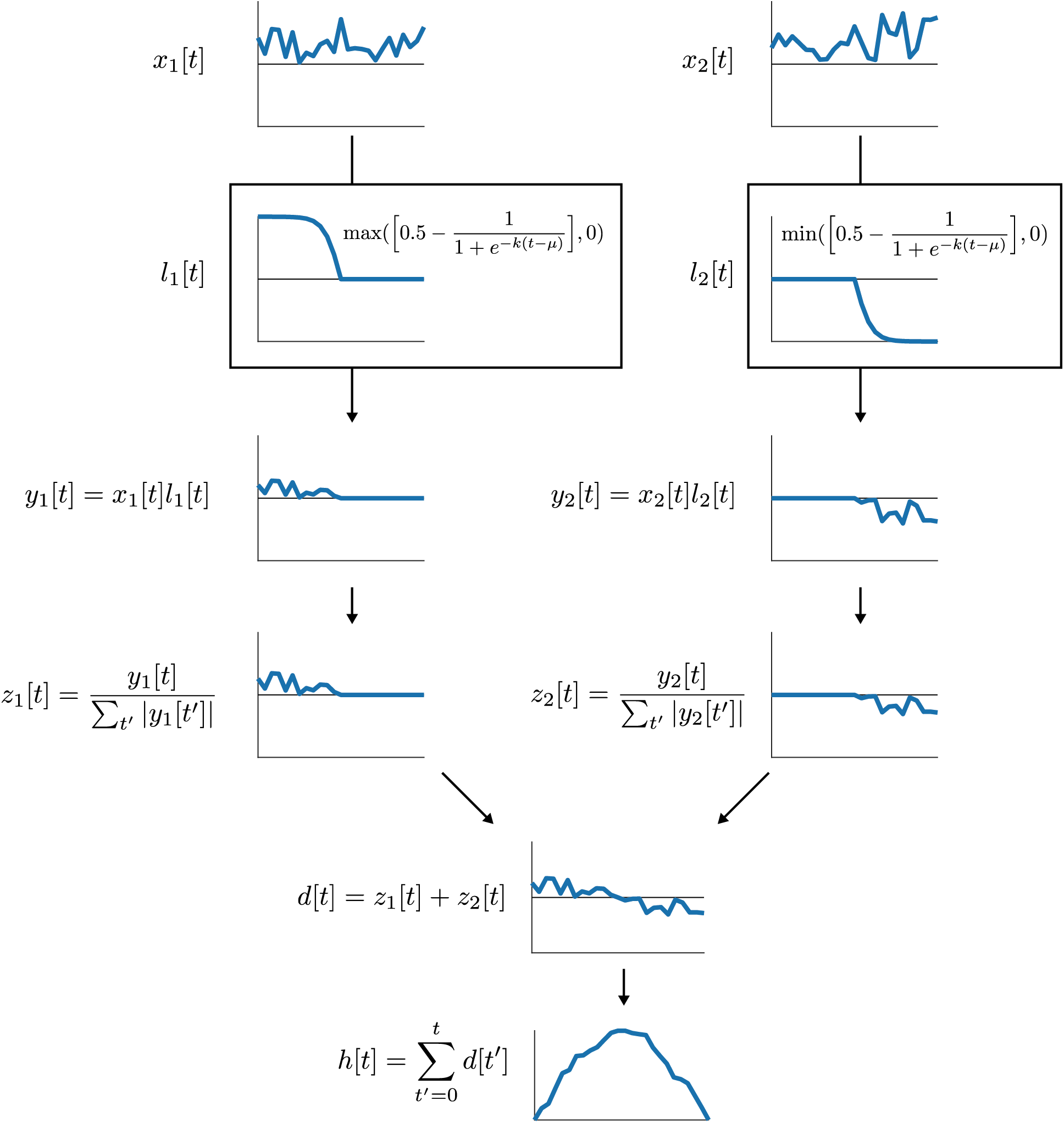
Constraining the smoothing kernel to be unimodal. This plot describes the set of operations (implemented in TensorFlow) that was used to constrain the smoothing kernel to be unimodal. Conceptually, the goal of these operations is to force the derivative to be exclusively positive for the first N time-points and then exclusively negative for the rest of the signal, thus preventing oscillations. We also must force the sum of the derivative to equal zero so that the kernel starts and ends at zero. Two positive vectors (themselves computed as the absolute value of real-valued vectors) were multiplied by a positively or negatively rectified logistic function with the same cross-over point. As a consequence, the first vector has positive values at the start of the signal, followed by zeros, and the second vector has negative values at the end of the signal, preceded by zeros. The two vectors were then normalized so that they sum to 1/-1. Finally, the two vectors were added and cumulatively summed, yielding a unimodal signal. The shape of the kernel is determined by the values of the two input vectors (*x*_1_ and *x*_2_) as well as the parameters of the logistic function (*μ* and *k*), all of which were learned. The input vectors were initialized with a vector of ones. *μ* was initialized to the value of the middle timepoint, and *k* was initialized to the value of 1 (and prevented from taking a value less than 0.001).

